# The MiDAS global genome catalog: 53,501 long-read MAGs representing all core prokaryotic genera in the global activated sludge microbiome

**DOI:** 10.64898/2026.07.10.737647

**Authors:** Lei Liu, Caitlin Margaret Singleton, Rasmus Hansen Kirkegaard, Mantas Sereika, Marie Riisgaard-Jensen, Kalinka Sand Knudsen, Aaron James Mussig, Jette Fischer Petersen, Zivile Kondrotaite, Miriam Peces, MiDAS Global Consortium, Philip Hugenholtz, Mads Albertsen, Per Halkjær Nielsen, Morten Kam Dahl Dueholm

## Abstract

Wastewater treatment relies on complex microbial communities, yet existing genome-resolved references for this essential engineered ecosystem remain dominated by short-read assemblies, limiting genome contiguity and linkage between taxonomic and metabolic function. We applied long-read sequencing to activated sludge from 83 globally distributed plants, reconstructing 53,501 metagenome-assembled genomes to establish the Microbial Database of Activated Sludge (MiDAS) global genome catalog. The catalog encompasses high-quality genomes for 12,047 prokaryotic species, 82% of which are not represented in GTDB release 226, and provides a median of 32 high-quality genomes for each of the 250 core prokaryotic genera previously defined in our MiDAS global 16S rRNA gene survey. This enables analyses of predicted functional traits and their ecological context, for example, we identified two sparsely represented *Nitrospiraceae* genera with conserved nitrite-oxidation genes that are abundant in higher-temperature wastewater treatment plants. In summary, the MiDAS genome catalog provides a framework for linking taxonomy, metabolism and ecological roles in wastewater treatment systems globally.

## Introduction

Diverse microbial communities drive activated sludge (AS) processes in wastewater treatment plants (WWTPs), and are responsible for nutrient removal (e.g., carbon, nitrogen, and phosphorus), protection of downstream habitats, and, increasingly, recovery of resources and energy^1^. Culture-independent molecular methods, particularly 16S rRNA gene amplicon sequencing and shotgun metagenomics, have been applied to these essential engineered ecosystems for decades and substantially advanced our understanding of WWTP taxa and community dynamics^2–6^. A recent large-scale metagenomic study has begun to provide genome-resolved references for global WWTP microbiomes^7^, but this resource is based on short-read assemblies, limiting genome contiguity and the linkage between taxonomic identity and metabolic function^8^.

The Microbial Database of Activated Sludge (MiDAS) project was developed to summarize knowledge about the physiology and ecology of functionally important microorganisms in activated sludge plants, anaerobic digesters, and related wastewater treatment systems^4,9^. This includes a 16S rRNA gene-based global survey of 740 WWTPs across 31 countries^10^. From this survey, we identified 965 ecologically important genera based on their relative abundance and ubiquity, with a subset of 250 core genera predicted to be central to AS functionality^10^. With the advent of high-throughput genome-resolved metagenomics^11–15^, it is now possible to extend MiDAS with a genome catalog, which we recently initiated using Danish WWTPs^16^. Long-read sequencing is particularly attractive for this purpose because it yields more contiguous metagenome-assembled genomes (MAGs), reduces the risk of contamination and mis-binning, and improves metabolic reconstructions^8^. Here, we extend our long-read MAG sequencing of AS ecosystems to 83 WWTPs across 27 countries to construct a comprehensive genome catalog of the global WWTP microbiome. This resource provides genome-resolved insights into the metabolic potential of core taxa, facilitates linking community composition to environmental and operational conditions, and reveals previously unrecognized nitrifying lineages that occasionally dominate nitrifier communities.

## Results

### Recovery of 53,501 MAGs from MiDAS global survey samples

To construct the MiDAS global genome catalog, we generated metagenomic datasets from 83 of 740 samples previously collected for the MiDAS global amplicon survey^10^. This subset is broadly representative of the survey encompassing 83 WWTPs across 75 cities, 27 countries, and five continents (**Fig. 1a**). Of these, 70 samples were from AS plants, 47 of which used advanced process configurations (**Fig. 1a and Supplementary Table S1)**. Each sample was sequenced using an entire PromethION flow cell (R10.4.1), yielding 11.3 Tbp of nanopore long reads >1 kbp. Reads had a median length of 6.1 kbp (IQR: 5.2-7.3 kbp), median read identity of 98.4% (Q18.0), and a median sequencing depth of 138 Gbp per sample (IQR: 127-155 Gbp) (**Fig. S1**).

**Fig. 1.**
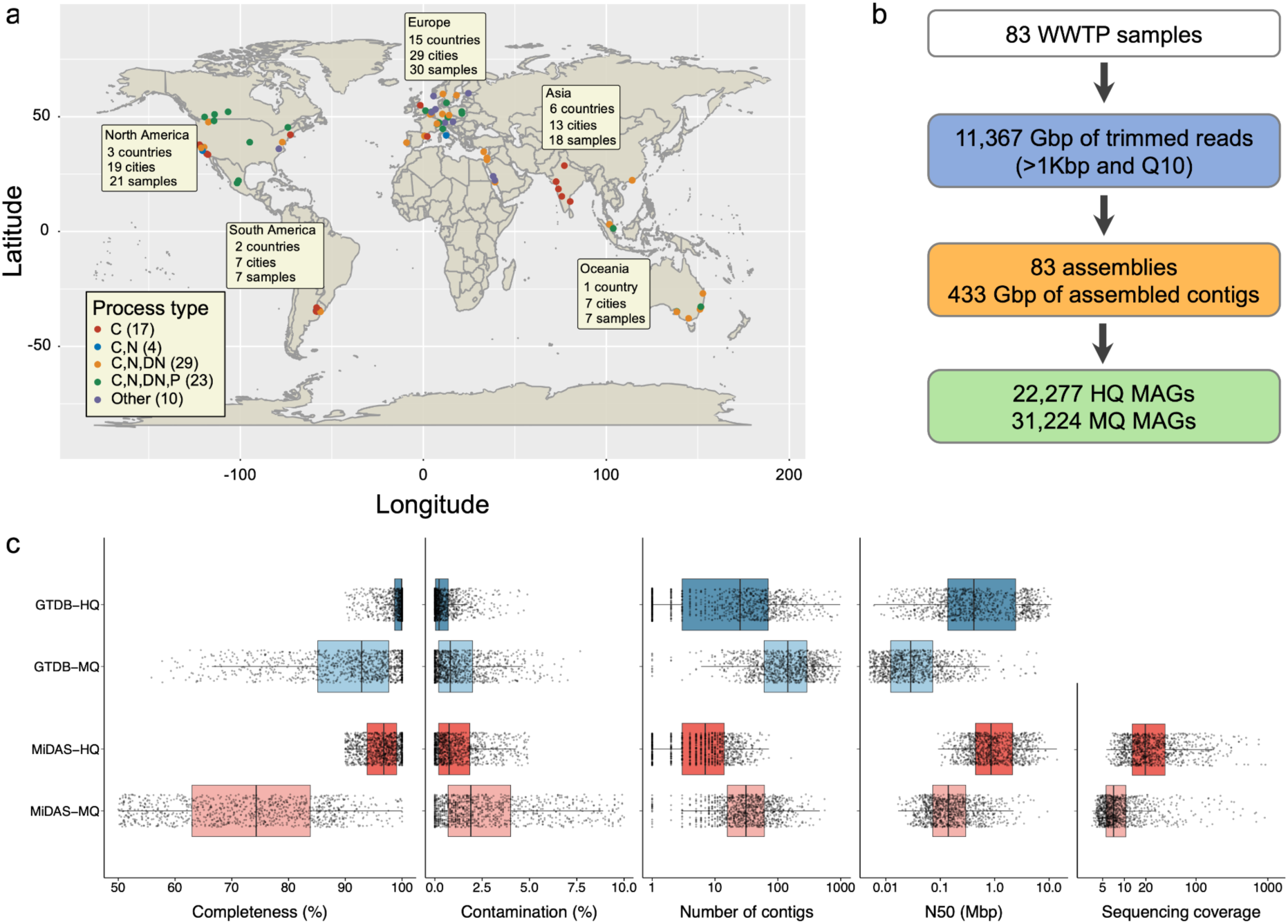
A vast high-quality resource of global WWTP microbiome data. **a,** Geographical and process-type distribution of MiDAS samples involved in this study. C indicates carbon removal, C,N indicates carbon removal with nitrification, C,N,DN indicates carbon removal with nitrification and denitrification, C,N,DN,P indicates carbon removal with nitrogen removal and enhanced biological phosphorus removal. **b**, Summary of primary data generated from MiDAS samples. **c**, Genome quality metrics of species representatives and comparison between MiDAS and GTDB databases using a subset of 1,000 randomly selected genomes to avoid visualization rendering challenges. HQ MAGs were defined using both CheckM and CheckM2 quality metrics and genome quality is shown as the higher of the two scores.

Each of the 83 datasets was assembled individually, producing 433 Gbp of contigs with a median read recruitment of 88.3% (IQR: 86.2-89.6%; **Supplementary Table S2**). Contigs from each sample were then binned, producing a total of 53,501 MAGs. These included 22,277 high-quality (HQ) MAGs with a median estimated completeness of 96.8% (IQR: 94.0-99.0%) and contamination of 0.9% (IQR: 0.3-2.0%; **Fig. 1b,c**).

HQ MAGs comprised a median of 9 contigs (IQR: 4-15) and a median N50 of 762 kbp (IQR: 421-1,686 kbp), indicating substantially lower fragmentation than those from other large-scale genome catalogs^13,14,17–25^ as well as HQ Genome Taxonomy Database (GTDB)^26^ reference genomes (**Fig. 1c; Supplementary Table S3**). Aligning with the MIMAG standard, all MiDAS HQ MAGs include complete sets of rRNA genes, which are often missed in short-read MAGs due to assembly and binning challenges^27^. This greatly facilitates linking the MiDAS global genome catalog to the 16S rRNA-gene based taxonomy and to numerous 16S rRNA-based AS surveys^16^.

HQ MAG sizes ranged from 0.53 to 14.8 Mbp (**Fig. S2**). The smallest MAG and other small MAGs are affiliated with known reduced phyla, including Patescibacteriota (median genome size: 0.99 Mb) and Iainarchaeota (median: 1.0 Mb)^28^. Whereas, the largest MAG and other large MAGs are affiliated with phyla known for more complex lifestyles, such as Myxococcota (median: 8.7 Mb) and Vulcanimicrobiota (8.4 Mb)^29^. Notably, 5.9% (1,308) of the HQ MAGs were identified as circular (cMAGs) by metaFlye^30^ and screened by *anvi-script-find-misassemblies*^31^ (**Fig. 2a**), thus representing the largest cMAG dataset to date^20,32,33^. In addition, 31,224 medium-quality (MQ) MAGs were recovered and included in the MiDAS global genome catalog. Species representatives derived from these MQ MAGs were still markedly less fragmented than the corresponding MQ reference genomes in GTDB, with ∼4.5-fold fewer contigs (32 versus 144) and ∼5-fold higher N50 (138 versus 27 kbp), highlighting the benefits of long-read assembly (**Fig. 1c**).

**Fig. 2.**
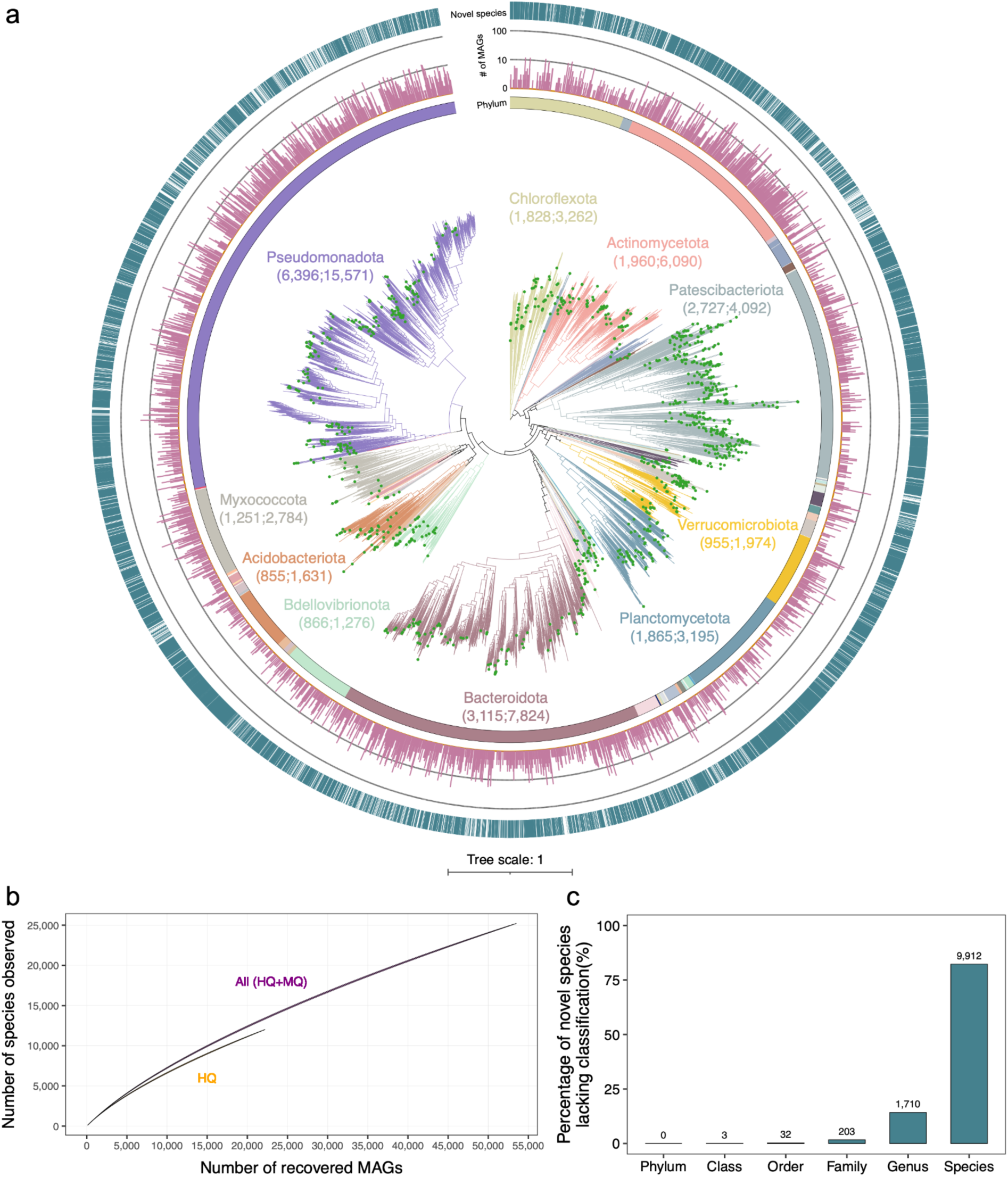
MiDAS global genome catalog populates extensive genomic diversity across the tree of life. **a**, Phylogenetic tree of 11,995 bacterial species representatives from the MiDAS-HQ catalog. The branches and inner strip chart are color-coded by phylum, with the 10 largest phyla displaying the number of species and MAGs in parentheses. The adjacent plot shows the number of HQ MAGs recovered for each species representative. The final strip chart shows species novelty based on the GTDB database, with colored strips indicating novel species. Green dots at the tips of the tree indicate circularized species representatives. **b,** Rarefaction curve depicting the number of HQ and total (HQ+MQ) species representatives assembled (y-axis) as a function of recovered MAGs (x-axis). **c,** Proportion and number of novel species lacking classification at different taxonomic levels based on GTDB (r226).

### Substantial unexplored and underdescribed lineages represented in the MiDAS global genome catalog

Genome dereplication clustered the 22,277 MiDAS HQ MAGs into 12,047 species representatives (11,995 bacterial and 52 archaeal). Relative to GTDB (r226), these species span 80 known phyla, 193 classes, 504 orders, 1,056 families, and 2,560 genera. Most species are novel (82%; 9,912), yet the species accumulation curve is far from saturation (**Fig. 2b**), consistent with estimates that the great majority of prokaryotic species remain unsequenced^34^. The distinct trajectory observed when MQ MAGs were included suggests that these genomes recover additional low-abundance diversity not captured among HQ genomes. Notably, 4,437 MQ-only species were recurrently detected across at least five samples and at least two non-source WWTPs while remaining below 1% maximum abundance.

To assess how much phylogenetic diversity MiDAS adds beyond rank-based novelty estimates, we integrated the catalog into a custom GTDB-derived HQ bacterial reference tree. Adding MiDAS HQ species increases total branch lengths by 26.0% and the number of HQ species representatives by 29.7% (**Fig. S4 and Supplementary Note**). This expansion reflected both improved representation of known taxa and the recovery of previously unrepresented lineages. Among known taxa, the catalog provides the first HQ species representatives for 3 phyla, 18 classes, 76 orders, 266 families, and 1,256 genera and at least doubles the number of HQ species diversity in a further 750 genera (**Fig. S3 and Supplementary Table S6**). Additionally, MiDAS adds HQ representatives for novel lineages at higher taxonomic ranks, including two classes, 21 orders, 104 families, and 1,040 genera that had no genomic representation in GTDB (**Fig. S5 and Supplementary Table S7**), revealing substantial previously unrepresented phylogenetic diversity within AS systems. Together, these results show that the MiDAS catalog expands both the species-level depth and the phylogenetic breadth of HQ genome-resolved references for AS systems.

### The MiDAS global genome catalog represents the global activated sludge microbiome

To quantify how well the MiDAS catalog represents global AS microbial communities, we first mapped reads from the 83 metagenomes used to construct the catalog back to MiDAS species-representative genomes and to the full non-dereplicated MiDAS genome catalog (**Fig. 3b and Fig. S6**). The MiDAS HQ species representatives captured a median of 57.6% of microbial reads (IQR: 52.9-61.2%), and inclusion of MQ species representatives increased median recruitment to 67.6% (IQR: 64.9-70.4%). The full non-dereplicated MiDAS catalog (n=53,501) further increased median read recruitment to 77.6% (IQR: 74.5-81.3%), where the increase suggests strain-level genomic diversity that may vary across WWTPs and indicates that the catalog captures most genomes detectable in the AS metagenomes generated in this study.

**Fig. 3.**
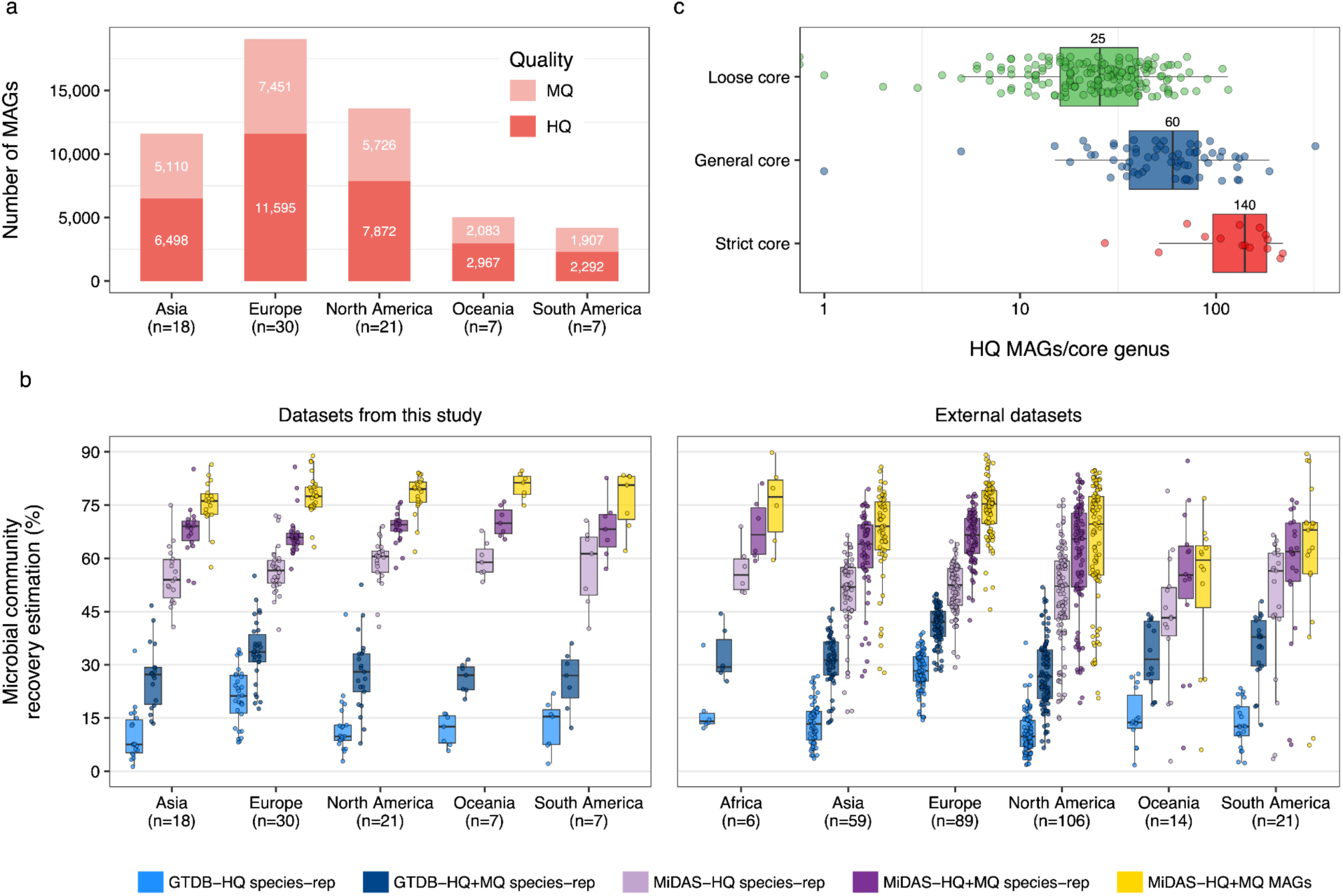
Towards a complete genome catalog for process-critical microbes in global WWTPs. **a,** Number of HQ and MQ MAGs recovered from different continents. n in the parentheses indicates the number of samples collected in each continent. **b**, Comparison of microbial community recovery across the 83 metagenomes analyzed in this study and 295 external datasets^16,36^ against different reference databases. The sets included GTDB HQ species representatives (n = 34,725), GTDB HQ+MQ species representatives (n = 136,646), HQ species representatives recovered from individual samples (n = 59-440), HQ+MQ species representatives recovered from individual samples (n = 166-1,105), MiDAS HQ species representatives (n = 12,047), MiDAS HQ+MQ species representatives (n = 25,162), and the full non-dereplicated MiDAS genome catalog (n = 53,501). Recovery ratio was estimated by mapping reads to the GTDB, MiDAS genome datasets in the left panel (Datasets from this study), with “species-rep” denoting the species representative dataset. Microbial community coverage was estimated using sylph in 295 independent, publicly available short-read activated sludge metagenomes not included in this study shown in the right panel (External datasets). **c**, Number of HQ MAGs per 16S rRNA gene-defined core genus in the MiDAS catalog. Core genera are classified as strict, general or loose based on relative abundance (>0.1%) and prevalence across WWTPs (strict >80%, general >50%, loose >20%; see Methods).

We then compared this representation with GTDB reference and sample-specific datasets. The GTDB HQ set provided limited coverage of most AS metagenomes (median: 13.0%, IQR: 8.0-19.7%). European samples showed relatively higher recruitment (median: 21.3%), which reflects the >1,000 HQ MAGs from Danish WWTPs^20^ included in GTDB (r226). By contrast, the MiDAS HQ catalog improved reads recruitment across regions, including underrepresented regions such as Oceania and South America, aligning with extensive sharing of species across AS systems^6,10^. HQ MAGs from individual samples explained only about one third of their own microbial communities (median: 37.8%, IQR: 28.9-44.3%) (**Fig. S6**).

Read-recruitment estimates closely matched coverage inferred using a computationally efficient heuristic *k*-mer-based approach implemented in sylph^35^ (**Supplementary Table S8**). Therefore, we applied sylph to estimate coverage for 295 publicly available AS metagenomes from 165 WWTPs across 17 countries and six continents^16,36^ (**Fig 3c**). The full non-dereplicated MiDAS catalog captured a median of 72.8% (IQR: 62.1-78.7%) of the microbial communities in these externally sourced, globally distributed datasets, further supporting its broad representation of global AS microbiomes.

### All activated sludge core taxa are represented in the MiDAS global genome catalog

Previously, we estimated that 250 core genera and 715 conditionally rare or abundant taxa (CRAT) genera are necessary for, or detrimental to, WWTP operation based on the MiDAS global amplicon survey^10^. All core genera are captured in the MiDAS genome catalog based on mapping of 16S rRNA genes using the genus-level identity threshold of ≥94.5%^37^. 248 of the 250 core genera (99.2%) are represented by at least one HQ MAG, with a median representation of 32 HQ genomes (IQR: 18-57) (**Fig. 3d**). ANI-based species clustering further resolved these into a median of 14 species per core genus (IQR: 7-23) (**Supplementary Table S9**). In total, 11,359 HQ MAGs (4,875 species) were assigned to core genera, accounting for more than half of all HQ MAGs in the MiDAS catalog. Across the 83 WWTP samples, these core genera contributed a median of 48.1% (IQR: 36.2-53.1%) of total community abundance, consistent with our previous amplicon-based estimate of 57-68% relative abundance across 589 AS samples^10^. Beyond the core genera, we recovered MAGs for 564 (78.9%) CRAT genera, including HQ MAGs for 509 (71.2%) genera with a median of 4 HQ MAGs each (IQR: 2-10). Taken together, the MiDAS catalog provides near-complete and genome-resolved coverage for core genera and CRAT lineages that structure global AS communities, thereby establishing a reference framework for future trait-based and process-level studies.

### Naming core AS species using HQ genome sequences as type material

Given the inferred ecological importance (prevalence and abundance) of AS core taxa to the performance of WWTPs, we decided to name all MAG-defined species (and associated higher ranks) currently lacking Latin names that belong to the 250 16S rRNA gene-defined core genera. This was done according to the SeqCode that uses genome sequences as type material^38^. For each genus, we selected a representative genome (MIMAG HQ, ≤10 contigs) by prioritizing circular MAGs over fragmented assemblies. In addition, geographic representation was incorporated during name assignment. Genus names were derived from the cities closest to the WWTPs from which type genomes were recovered, while species epithets were assigned as generic, WWTP-related or geographic names to minimize redundancy (**Method and Supplementary Table S10**). This provided names for 1,474 species, 140 genera, 41 families, 15 orders and 4 classes, which would support scientific communication about these important AS taxa.

### Conservation of inferred functional traits in genomically well-represented core genera

In AS ecosystems, genus is often considered an ecologically meaningful unit because closely related species within a genus frequently share overlapping functional roles^10,39^. The large number of independent MAGs recovered for most core genera enabled us to directly test this assumption and quantify how consistently genome-encoded functional traits are conserved within genera. To obtain robust, genome-based trait estimates, we used GTDB-defined genus-level groups rather than 16S rRNA gene-based genus definitions. This reclassification yielded 277 core genera with at least 10 HQ MAGs (**Fig. 4**), reflecting the different approaches to define taxa by SILVA (16S rRNA gene sequence identity thresholds^37^) and GTDB (relative evolutionary divergence normalization^40^). We classified each genome-encoded trait as either conserved present (when detected in ≥80% of MAGs within a genus), conserved absent (<20% of MAGs in a genus) or mixed (20-80% of MAGs in a genus).

**Fig. 4.**
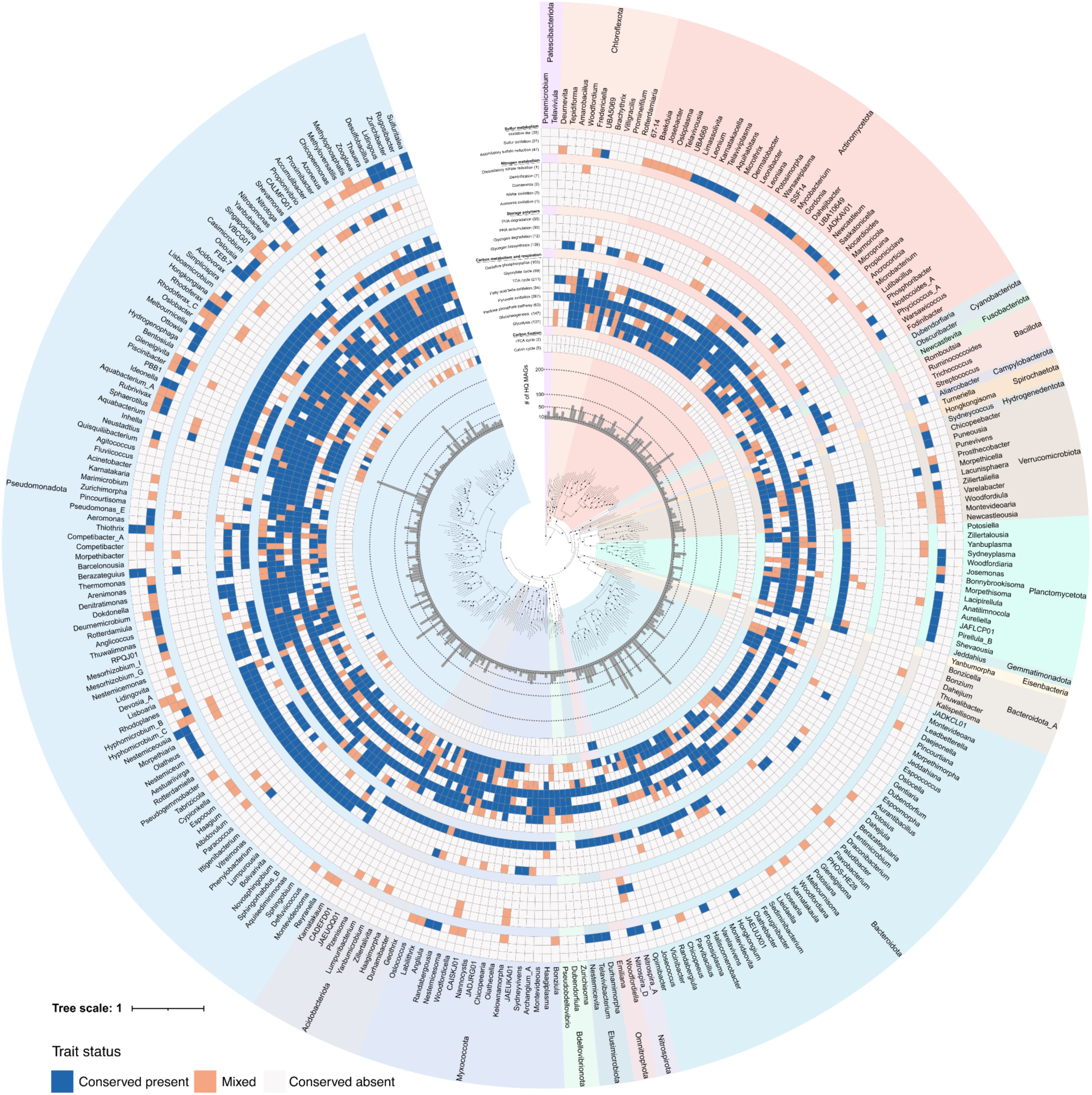
Metabolic trait profiles of the 277 core genera identified in global WWTPs. The inner ring indicates the number of HQ MAGs recovered for each core genus. Heatmap rings illustrate the percentage of HQ MAGs within each core genus that encode a given metabolic pathway. Only pathways with complete (100%) gene representation in the KEGG or customized modules were considered present. Traits were classified as conserved present when a complete trait was detected in ≥80% of MAGs, conserved absent when detected in <20% of MAGs, or mixed when detected in 20-80% of MAGs (The exact percentages are available in **Supplementary Table S13**). The outer rings display the names of core genera along with their affiliated phyla.

Across the selected 6,094 genus-trait combinations examined, 24.3% were conserved present, 67.3% were conserved absent, whereas only 8.4% were mixed (**Fig. 4 and Fig. S7a**), indicating substantial functional consistency in trait presence or absence within core genera. However, this consistency was pathway-dependent (**Fig. S7b**). Mixed states were rare for traits such as nitrite oxidation, the reductive tricarboxylic acid cycle and pyruvate oxidation, but exceeded 20% for glycolysis, the pentose phosphate pathway and assimilatory sulfate reduction. Thus, genus-level groupings provide a useful but pathway-dependent framework for functional interpretation.

The strongest conservation was observed for carbon metabolism. Complete pyruvate oxidation and the tricarboxylic acid (TCA) cycle were detected in nearly all core genera (272 and 259 genera) and were classified as conserved present in 267 (96%) and 211 (76%) genera, respectively. By contrast, autotrophic carbon fixation capacity was rare and restricted to a few specialized lineages. The Calvin cycle was classified as conserved present in five genera, including *Sulfuritalea*, whereas the reductive TCA (rTCA) cycle was conserved in two genera, including *Nitrospira*_A *and Nitrospira*_D. Complete oxidative phosphorylation potential was encoded in 103 core genera, of which 75 (27%) were classified as conserved present, likely reflecting the temporal redox conditions occurring in AS that support metabolic niche differentiation (see below).

Intracellular storage compounds are often used by bacteria experiencing fluctuating substrate availability^41^. Accordingly, glycogen and polyhydroxyalkanoates (PHA) were broadly represented across the core community, with 230 genera encoding biosynthetic potential for at least one of the two polymers and 192 genera (69%) showing conserved present storage potential. Glycogen biosynthesis was detected in 191 genera and classified as conserved present in 139 genera, whereas PHA biosynthesis was detected in 113 genera and conserved in 90 genera. However, complete storage/mobilization potential was far more common for PHA than for glycogen, with only 8 genera encoding both glycogen biosynthesis and degradation potential, compared with 62 genera for PHA (**Fig. 4)**. This observation suggests that PHA-utilizing lineages more commonly retain pathways for both intracellular carbon storage and mobilization, potentially coupling them to short-term fluctuations in carbon availability and redox conditions^42^, although incomplete annotation of glycogen degradation genes may also contribute to this difference.

As expected, nitrification potential was confined to a narrow and highly specialized guild within the core community^10^. Ammonia oxidation potential occurred only in the canonical ammonia-oxidizing bacteria (AOB) genus *Nitrosomonas* (100% conservation) and in the comammox genus *Nitrospira*_D (77% conservation), according to the GTDB taxonomy. Conserved nitrite-oxidation potential was largely restricted to *Nitrospira*_D (100% conservation), *Nitrospira*_A (96%), and the canonical NOB genus *Nitrotoga* (100%). By contrast, denitrification traits were broadly distributed across the core community. Only seven genera (2.5%) were classified as conserved present for complete denitrification pathway, however, 215 genera (78%) showed conserved potential for at least one reduction step (nitrate, nitrite, nitric oxide, or nitrous oxide reduction). This pattern is consistent with denitrification being modular and distributed across multiple taxa, rather than concentrated in a small set of complete denitrifiers^43^. Notably, 69 genera (25%) encoded conserved nitrous oxide reduction potential (*nosZ*), including two genera that lacked a complete upstream denitrification pathway, forming a widespread guild of putative N_2_O reducers that may act as community-level N_2_O sinks (**Fig. S8**). By predicting which core lineages encode complete versus partial nitrogen-transformation pathways, the catalog provides a taxonomically anchored framework for interpreting nitrogen cycling and pinpointing putative N_2_O-consuming guilds in wastewater treatment systems.

Although sulfur cycling is not generally considered a major process in AS^10,36^, the sulfur oxidation (Sox) module was classified as conserved present in 27 genera (10%). In addition, oxidative dissimilatory sulfate reduction (Dsr) capacity, represented by phylogenetically confirmed reverse-acting *dsrAB* genes, was conserved in 11 genera (4%). This pattern contrasts with the restricted nitrifying guild and suggests that sulfur-oxidation capacity is more broadly embedded within the core community. Together, these results show that conserved-present traits are widespread in carbon metabolism (99% of genera), denitrification (78%), carbon storage (69%) and, to a lesser extent, sulfur oxidation (10%), whereas carbon fixation (2.5%) and nitrification (2%) are highly restricted traits.

### Two additional *Nitrospiraceae* genera expand the diversity of nitrite oxidizers

Functional trait analysis showed that nitrite oxidation potential was primarily associated with classical NOB, while examination of the entire set of HQ MAGs identified two additional *Nitrospiraceae* genera encoding conserved nitrite oxidation genes. These genera, here proposed as *Nitrosoma* gen. nov. (JAHBWI01 in GTDB r226) and *Nitrocyclum* gen. nov. (BQWY01 in GTDB r226), were each represented by only two public MAGs before the addition of MiDAS genomes. Public MAGs now assigned to *Nitrocyclum* had previously been described as *Nitrospira* or *Nitrospiraceae* nitrite oxidizers from brackish landfill leachate treatment^44^ and deep-underground saline spring biofilms^45^ rather than BQWY01. MiDAS *Nitrocyclum* genomes represent distinct novel species-level members recovered from AS. By contrast, two public MAGs now assigned to JAHBWI01 recovered from AS^46^ and soil^13^ had not, to our knowledge, been previously discussed as representatives of a putative nitrite-oxidizing lineage.

All recovered members of *Nitrosoma* gen. nov. (6 HQ MAGs representing 1 novel and 1 GTDB-represented species) and *Nitrocyclum* gen. nov. (7 HQ MAGs representing 5 novel species) (**Supplementary Table S10**), encode up to four complete copies of *Nitrospira*-like *nxrAB* (**Fig. 5a**). This is in contrast to other *Nitrospiraceae* genera that usually encode 1-2 copies of *nxrAB*, which may support higher production of nitrite oxidoreductase and/or substrate affinity specialization^47^ in *Nitrosoma* and to a lesser extent *Nitrocyclum*. The *nxrA* gene tree indicates that the four copies of this gene in each member of the genus *Nitrosoma* arose through three successive duplications, the first in the *Nitrospiraceae* ancestor and the last following speciation (**Fig. 5a**). In addition, both genera encode key genes for carbon fixation *via* the rTCA cycle, consistent with the autotrophic lifestyle typical of *Nitrospiraceae* (**Figure S9**).

**Fig 5.**
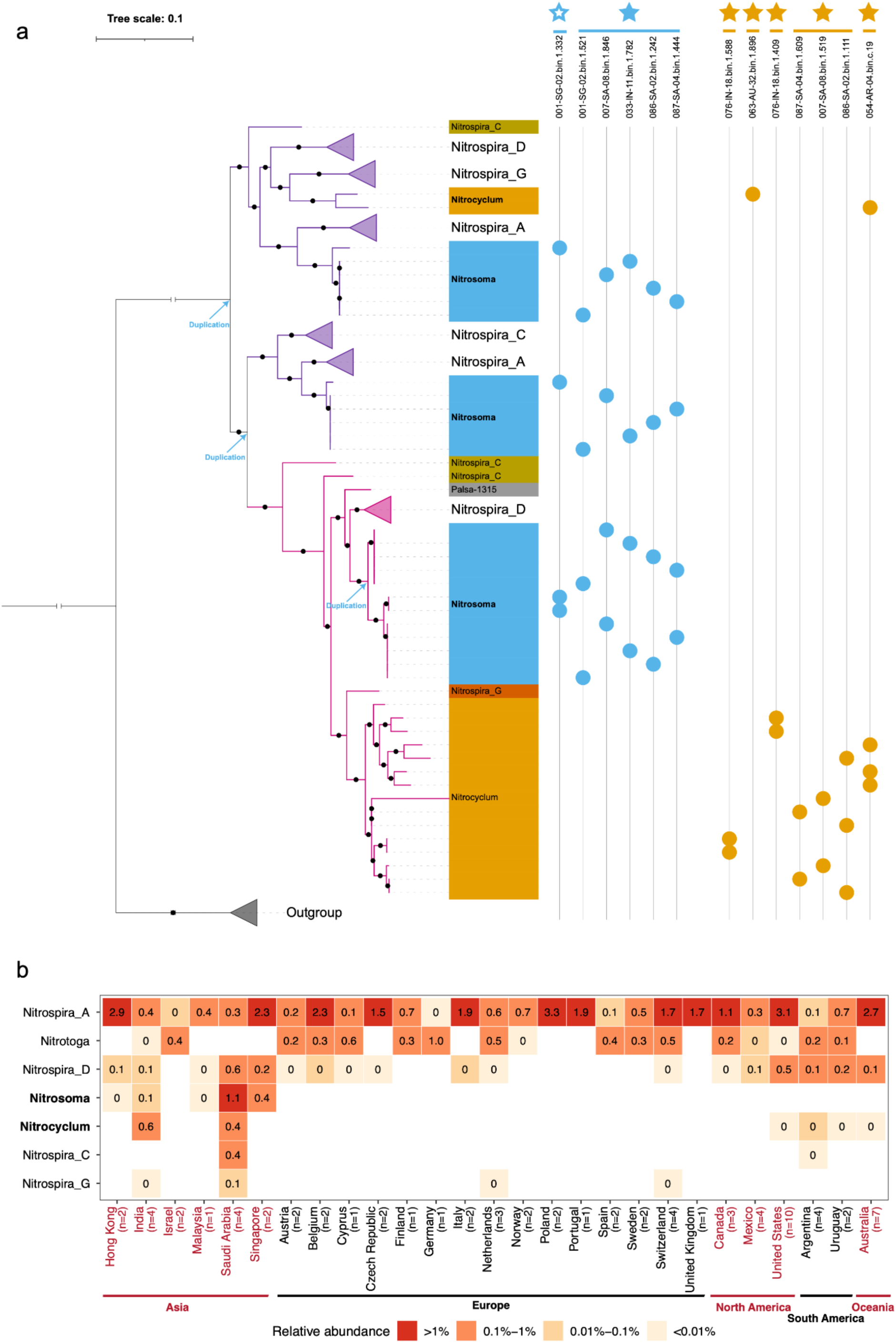
*NxrA* phylogeny and global distribution of sparsely represented *Nitrospiraceae* nitrite oxidizers. **a,** Maximum-likelihood phylogeny of *nxrA* gene sequences from *Nitrosoma* gen. nov. and *Nitrocyclum* gen. nov. together with selected representative nitrite-oxidizing lineages. Collapsed triangles indicate reference clades and the outgroup (*Nitrotoga*), black circles indicate nodes with bootstrap support >70%, and arrows mark inferred NxrA duplication events. The purple and red clades were assigned to *Nitrospira* clade 1 and *Nitrospira* clade 2, respectively, using GraftM. The filled stars indicate the novel species, whereas open stars indicate species that have been represented in GTDB (r226). Coloured blocks denote genus-level groups, and the matrix to the right links *nxrA* gene copies to their source MiDAS HQ MAGs. Each horizontal line indicates a species-level group, which may include one or multiple MAGs. Blue and orange symbols indicate *Nitrosoma* and *Nitrocyclum* genomes, respectively. **b,** Country-level distribution of major nitrite-oxidizing genera across global WWTP metagenomes in this study. Cell colours and values show the mean genus-level relative abundance across metagenomes from each country or region. Values are rounded to one decimal place, and blank cells indicate no detection.

We evaluated 378 globally distributed metagenomes^16,36^ and detected both genera in more than 20 WWTPs across multiple countries and continents, indicating a geographically widespread yet infrequent occurrence. Notably, neither genus was detected in European WWTP samples despite this being the most well represented region in our survey (**Fig. 5b**). Consistent with this pattern, both genera were preferentially detected in tropical and subtropical regions with WWTPs operating at higher-temperatures (∼30 °C vs ∼20 °C; Wilcoxon test, *P* < 0.01). In some instances, these genera reached high relative abundances, e.g., up to 2.4% *Nitrocyclum* in an Indian WWTP and 3.2% *Nitrosoma* in a Saudi Arabian WWTP (**Figure S10**). No consistent associations between relative abundance and process configuration or plant type were observed, suggesting that temperature, together with local factors such as influent composition^48–50^ and plant-specific operating conditions, may jointly influence the occurrence and abundance of these genera. Overall, canonical NOB dominated most WWTPs, whereas *Nitrosoma* and *Nitrocyclum* were infrequent but occasionally exceeded total canonical NOB abundance in higher-temperature systems (**Fig. 5b**).

### Species-level differentiation of polyphosphate-accumulating organisms

Given the central importance of enhanced biological phosphorus removal (EBPR) in wastewater treatment systems, we next examined the genomic diversity and functional differentiation of polyphosphate-accumulating organisms (PAOs). Here we demonstrate the ability to perform species-resolved comparisons enabled by the large number of HQ MAGs in the MiDAS catalog. We focused on species representatives of four experimentally verified PAO genera; *Ca.* Accumulibacter, *Azonexus*, *Ca.* Phosphoribacter and *Ca.* Lutibacillus^51–53^ (**Fig. 6 and Supplementary Table S11**).

**Fig. 6.**
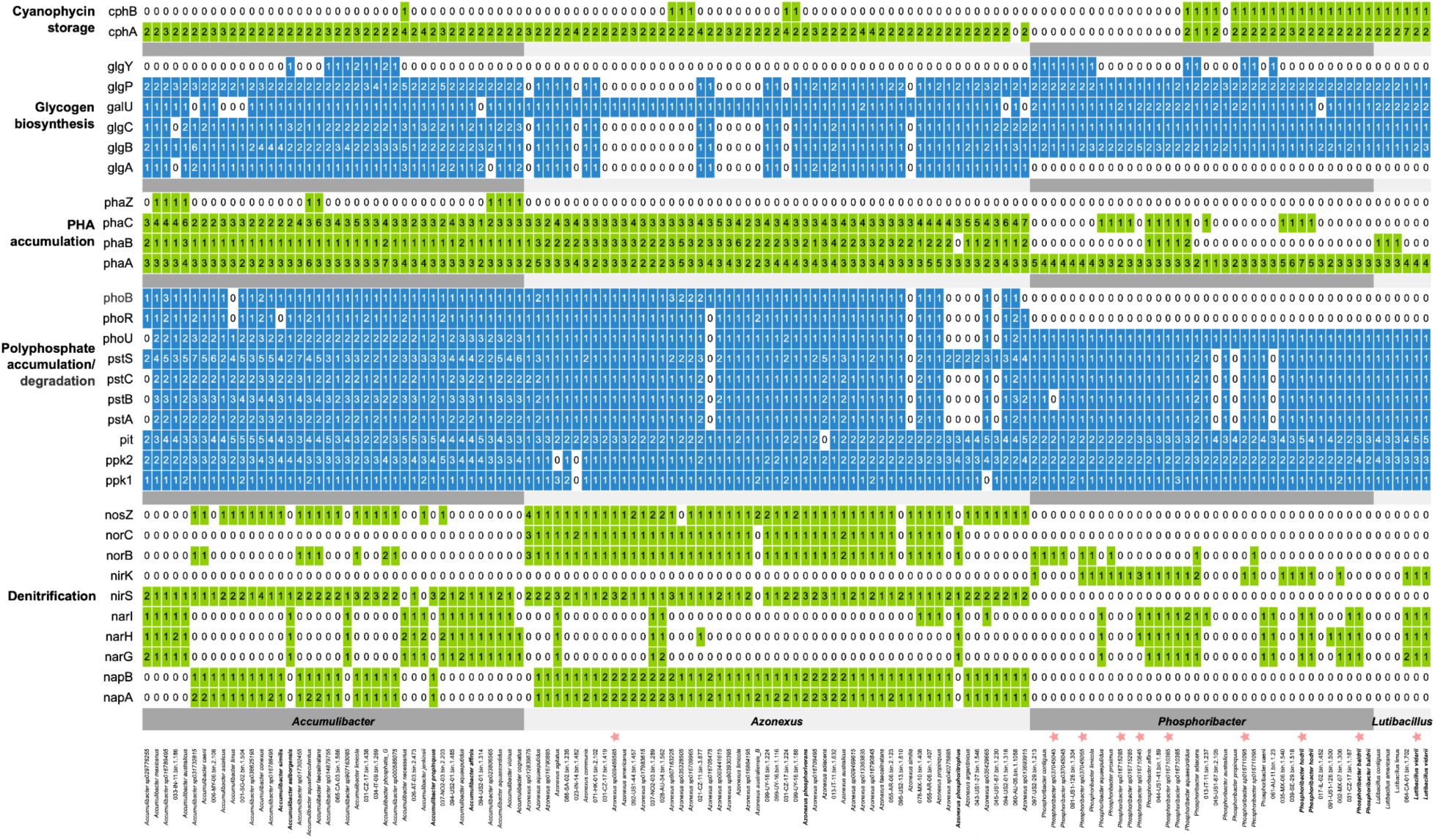
Prevalence and distribution of key genes across the four experimentally verified PAO genera based on HQ species genomes from the MiDAS and GTDB catalogs. HQ MAGs from GTDB (r226) are marked with orange stars. The numbers within each cell represent the copy number of the corresponding gene identified in each MAG.

These PAOs thrive in anaerobic-aerobic systems by using inorganic polyphosphate and organic polymers as dynamic storage compounds^54,55^ (**Supplementary note**). All species representatives encode systems for phosphate transport and polyphosphate accumulation with lineage-specific patterns for metabolism of organic storage polymers. Consistent with PHA being a major organic storage polymer in PAOs, PHA biosynthesis and degradation genes were universally detected in *Ca.* Accumulibacter and *Azonexus*. Notably, five *Ca*. Phosphoribacter species also encode *phaABC*, suggesting potential PHA-based storage in this genus, even though this trait is not generally reported for *Ca.* Phosphoribacter^55^. Glycogen represents another key organic storage polymer in *Ca*. Accumulibacter and *Azonexus*^55^. However, a distinct subset of *Azonexus* species lacked key genes (*glgABC* and *glgP*), indicating that not all members of this genus use the well-described glycogen-based storage strategy, or that they rely on alternative, non-canonical enzymes that remain to be identified. In addition, the capacity to synthesize and degrade cyanophycin, a potential nitrogen-rich intracellular storage polymer^53^ (not specific to PAOs), was supported by the presence of both biosynthesis and degradation genes. These genes were uniformly detected in *Ca.* Lutibacillus species, whereas in *Ca.* Phosphoribacter they were restricted to a phylogenetically clustered subset of species (16/28). Collectively, these patterns reveal the functional diversification of storage polymer metabolism among PAO lineages, with different genera, and even closely related species, encoding distinct combinations of glycogen, PHA, and cyanophycin alongside polyphosphate accumulation. Whether all species belonging to these four genera have the PAO phenotype remains to be experimentally investigated.

Denitrification potential, an important trait in EBPR-based biological nutrient removal systems^56^, further differentiated species belonging to the four PAO genera. In *Azonexus*, most species (77%) encode a complete denitrification pathway, forming a dominant PAO-denitrifier ecotype in which polyphosphate metabolism is consistently coupled to the full potential of nitrate-to-dinitrogen reduction. For *Ca.* Accumulibacter, almost all species (93%) encode NirS-type nitrite reduction but lack NorBC for further conversion of NO. For nitrate and nitrous oxide (N_2_O) reduction, two mutually exclusive ecotypes were observed (**Figure 6**). One (21/40 species) encodes periplasmic NapAB and NosZ, consistent with the capacity for nitrate-to-nitrite reduction under nitrate-limited conditions and N_2_O to N_2_, respectively. The other (18/40) lacks NosZ and encodes the membrane-bound NarGHI, which is generally associated with nitrate-to-nitrite reduction under nitrate-replete anoxic conditions^57,58^. This partitioning suggests distinct nitrate-reduction strategies among *Ca.* Accumulibacter species under fluctuating redox conditions^57,59^. In *Ca*. Phosphoribacter and *Ca.* Lutibacillus, nitrate reduction potential was present in a subset of species, with NarGHI detected in 16 of 33 species and frequently co-occurring with NirK (12/16). By contrast, 13 species lacked detectable denitrification genes or complexes. Additionally, these species-level patterns were further supported by high within-species consistency of PAO-related traits: 95.8% of species-trait combinations were conserved within PAO species represented by at least three HQ MAGs, and 95.5% of genus-level mixed traits were classified as conserved states at the species level. Taken together, these results indicate that each PAO genus comprises multiple species-level ecotypes, suggesting that niche partitioning in EBPR systems occurs not only among PAO genera but also among closely related species within the same lineage.

## Discussion

A key question in genome-resolved microbial ecology is how functional potential is organized across taxonomic scales in complex communities. Across core genera in the global AS microbiome, our results reveal a scale-dependent organization in which some traits reflect lineage-level constraint, others are partitioned among closely related species, and modular pathways are assembled across multiple taxa. This provides a framework for interpreting AS functions as traits shaped by different degrees of evolutionary conservation and ecological partitioning, rather than as uniform properties of genus-level taxa. Carbon metabolism formed an expected broadly distributed chemoorganoheterotrophic scaffold, suggesting that presence or absence of these basic modules alone provides limited resolution for distinguishing ecological roles among core taxa. By contrast, process-relevant traits showed stronger differences in their taxonomic organization, indicating that functional interpretation requires attention to the biological scale at which each pathway is conserved or distributed. However, these trait predictions remain limited by database-dependent annotation biases, which may lead to under-detection of divergent or poorly characterized genes.

Consistent with the established understanding of nitrifiers, nitrification potential was restricted to specialized taxa. However, long-read genome analysis uncovered diversity within this constrained functional guild. Comammox bacteria, whose identity cannot be reliably resolved from 16S rRNA gene data alone, were represented by 13 HQ MAGs (10 species) including 4 novel species, supporting their broader presence in full-scale WWTPs^60^. Moreover, the identification of *Nitrosoma* gen. nov. and *Nitrocyclum* gen. nov. expands the known diversity of putative NOB in WWTPs. For *Nitrocyclum*, the MiDAS catalog extends prior observations from individual non-AS MAGs by resolving distinct novel species-level members in AS and linking them to conserved *nxrA* across multiple WWTPs. For *Nitrosoma* gen. nov., the MiDAS genomes newly link this previously functionally unresolved lineage to nitrite oxidation potential. Notably, this genus consistently retains multiple distinct phylogenetic clades of the *nxrA* gene, suggesting stable retention of functionally divergent Nxr variants within the same lineage. Such genomic configurations may enable nitrite oxidation across a broader range of redox or substrate conditions. However, the catalytic properties, regulation or environmental responsiveness of these populations carrying distinct Nxr enzymes will require further experimental assessment.

Denitrification potential represents a contrasting mode of functional organization. Although denitrification is well recognized as a modular pathway in different ecosystems^43,61^, our analysis places this modularity into a global taxonomic context by showing how individual pathway steps are distributed across recurrent core genera. Thus, genus-level profiles can inform denitrification potential only when linked to step-specific genome annotations. Within this modular process, the widespread canonical NosZ*-*encoding guild identified here suggests substantial potential for N_2_O reduction and may represent a target for N_2_O emission mitigation in WWTPs^62^. In addition, species encoding the recently described L-NosZ variant^63^ were detected in 28 of 83 WWTPs, but their combined relative abundance remained low where present (median 0.004%, IQR 0.002-0.013%; maximum 0.3%). This suggests that L-NosZ-encoding populations are unlikely to dominate community-level N_2_O reduction in the system, although low relative abundance does not necessarily preclude ecological importance^64^.

Species-level analyses of four experimentally verified PAO genera revealed another layer of functional organization. Rather than following a single conserved genomic configuration, closely related PAO species differed in both storage strategies and nitrate-reduction potential^55^. The expanded genomic representation provided by the MiDAS catalog resolves this variation across major PAO lineages, including Nap-based complete denitrifiers in *Azonexus*, coexisting Nap- and Nar-based lineages in *Ca.* Accumulibacter, and Nar-dominated, truncated denitrifiers in *Ca.* Phosphoribacter and *Ca.* Lutibacillus. This species-level mosaic is consistent with PAO niche partitioning across microscale chemical gradients within flocs and the alternating anaerobic, anoxic and aerobic zones of EBPR systems, where oxygen and nitrate availability vary over space and time^65^. Together, this species-level resolution complements genus-level trait patterns observed across the core community (**Supplementary Note**) and suggests that EBPR performance and robustness may depend not only on total PAO genus abundance but also on the relative contributions of distinct species adapted to different microscale environments. More broadly, the quality of the MiDAS catalog enables analogous species-resolved analyses for other functionally important guilds, such as nitrifiers and key hydrolytic lineages^66^, linking microbial diversity to process conditions and performance.

Overall, the MiDAS genome catalog substantially expands microbial representation and metagenomic resolution for global WWTPs. Its high contiguity makes it particularly valuable for biological questions that are difficult to resolve with short-read MAGs alone, including linking full-length rRNA genes to genome-encoded traits, resolving repeated functional genes, and preserving complete operon, biosynthetic gene cluster and mobile element context. These features support identification of novel functional guilds^16,53^, analyses of antibiotic resistance genes^67^ and host-virus/plasmid linkages^68,69^, primarily in engineered wastewater treatment systems and potentially in related environments. Together with near-complete gene inventories, the catalog also provides a foundation for metagenome-guided cultivation^70–73^, including auxotrophic predictions^74,75^ to target uncultured microorganisms and experimentally test genome-derived hypotheses.

## Methods

### Sample collection, DNA extraction, and long-read sequencing

The samples included in this study were primarily collected in 2018 for the MiDAS 4 study^10^, except for one AS sample collected in 2023 (**Supplementary Table S1**). These samples were contributed by the MiDAS Global Consortium, which is distributed across various regions worldwide. The sequencing and analysis of all 83 samples were approved by the relevant authorities, either through applications under the Nagoya Protocol or by authorization from local WWTP managers. Detailed sampling procedures have been described previously^10^. To ensure DNA quality, samples stored at -80 °C were used for fresh DNA extraction using the DNeasy PowerSoil Pro Kit (QIAGEN, Germany), following the manufacturer’s instructions. Bead-beating was performed using vortexing at 80% of maximum speed to ensure efficient lysis while minimizing excessive DNA shearing. DNA was quantified using a Qubit 4 Fluorometer (Thermo Fisher Scientific, MA, USA).

To enable efficient long-read sequencing, 2.5 μg of DNA per sample underwent one round of size selection using the long fragment buffer to enrich fragments >3 kbp before library preparation. The concentration and quality of size-selected DNA were re-evaluated using the Nanodrop ND1000 (Thermo Fisher Scientific, MA, USA) and Qubit 4 Fluorometer (Thermo Fisher Scientific, MA, USA). Libraries were prepared for each sample using the Ligation Sequencing DNA V14 (SQK-LSK114) protocol, then loaded onto R10.4.1 flow cells (FLO-PRO114M) and sequenced using a PromethION P24 device, with one sample per flow cell. The kit 14 chemistry was run at a 4 kHz sampling rate for phase 1 samples and later at 5 kHz for phase 2 samples (**Fig. S1** and **Supplementary Table S2**). Basecalling was performed with Guppy (v6.3.9-v6.5.7) integrated into MinKNOW, using the super-accurate model with a Phred score of >10. The basecalled FASTQ files were further filtered to remove reads shorter than 1 kbp using seqkit (v2.6.1)^76^ to generate trimmed long reads. Statistical summary (such as mean, median read length, read quality, and N50) for each long-read dataset was calculated by NanoStat^77^ (v1.6.0). GNU Parallel (20210822)^78^ was used to execute tasks in parallel.

### MAG recovery, quality assessment, taxonomic assignment, and species clustering

The genome recovery pipeline mmlong2 (v0.9.2; https://github.com/Serka-M/mmlong2)^15^ with the parameter ‘*-sem wastewater*’ was used to recover MAGs from each of the 83 samples individually. Briefly, clean reads were first assembled into contigs using MetaFlye (v2.9.2)^30^, followed by one round of polishing with Medaka (v1.8.0; https://github.com/nanoporetech/medaka). Eukaryotic sequences identified by Tiara (v1.0.3)^79^ were removed before performing four rounds of iterative genome binning with three binning tools: MetaBAT2 (v2.15)^80^, SemiBin (v1.5)^81^, and GraphMB (v0.1.5)^82^. Contigs with lengths < 3 kbp were discarded. DAS_Tool (v1.1.3)^83^ was then used to select the best representative bins from the binning iterations for each metagenome. GUNC (v1.0.5)^84^ was used to detect the chimerism and contamination in recovered MAGs. Genome quality assessment and coverage estimation within each sample were conducted using CheckM2 (v1.0.2)^85^ and CoverM (v0.6.1)^86^. Only MAGs that met at least the minimum MQ criteria were retained in the MiDAS catalog. HQ MAGs were evaluated according to the MIMAG guidelines^87^, with rRNA genes identified using Bakta (v1.8.1)^88^ and Barrnap (v0.9; https://github.com/tseemann/barrnap), while tRNA genes were detected with tRNAscan-SE (v2.0.11)^89^. Additionally, 16S rRNA genes were annotated using SILVA (v138)^90^ and the MiDAS database (v5.0) with VSEARCH (v2.22.1)^91^.

To ensure the latest classification, after the default run of mmlong2, we re-run the classification steps using GTDB-Tk (v2.4.1)^92^ with GTDB^26^ (r226) and the MiDAS database (v5.3)^93^. We also re-run the classification based on the MiDAS database (v4.8) to have a consistent group with the previous study^10^, where core taxa and CRAT were identified. In this study, two independent methodologies, CheckM (v1.2.2; lineage-specific workflow)^94^ and CheckM2, were employed to estimate genome completeness and contamination, ensuring robust quality assessment and minimizing potential biases associated with specific lineages. Based on these results, 22,277 out of 53,501 MAGs were classified as HQ according to the stringent MIMAG standards, while the remaining were designated as MQ. The 53,501 recovered MAGs form the full non-dereplicated MiDAS global genome catalog (MiDAS-HQ+MQ), with a subset of 22,277 HQ MAGs constituting the MiDAS-HQ catalog. To identify distinct species within the MiDAS-HQ+MQ and MiDAS-HQ catalogs, dRep (v3.5.0)^95^ was used to cluster the two genome sets at 95% average nucleotide identity (ANI) with the parameter of *‘dereplicate’* and ‘*--S_algorithm skani*’. Additionally, MAGs within each species cluster were extracted from dRep output results. Notably, HQ species representatives in GTDB (r226) were selected using the same criteria.

### Phylogeny tree construction

The phylogeny of 11,995 bacterial HQ species representatives within the MiDAS catalog was inferred using GTDB-Tk (v2.4.1) with the *de_novo_wf* workflow, based on 120 bacterial marker genes. This process utilized dependencies including Prodigal (v2.6.3)^96^, pplacer (v1.1.alpha19)^97^, FastTree 2 (v2.1.11)^98^, skani (v0.2.1)^99^, and HMMER (v3.4)^100^. The resulting tree was then imported into iTOL (v6)^101^ for further refinement and visualization.

### Microbial community coverage estimation

We randomly selected 1% of reads from each metagenome using seqkit (v2.6.1), with a median size of 1.4 Gbp (IQR: 1.3-1.5 Gbp), to estimate the percentage of the microbial community that was successfully recovered and to reduce the computational cost of mapping all metagenomes. The subsampled reads were then mapped to the different MiDAS genome sets in Fig. 3b (Dataset from this study) and Fig. S6 using minimap2 (v2.26-r1175)^102^ and filtered with the following criteria: a minimum alignment length of 1 kbp, a minimum identity of 90%, and a minimum read coverage of 50%. For datasets that include only species representatives, we also used a *k*-mer-based method, sylph (v0.8.0)^35^, to calculate the known percentage based on the whole metagenomes. This approach also showed high consistency with the minimap2-based mapping method (**Supplementary Table S8**). The coverage estimation of GlobDB^103^, a more comprehensive species-dereplicated microbial genome resource that integrates additional genome catalogues beyond GTDB, was also included for the comparison. For publicly available, external 295 datasets, sylph was used to calculate the recovery fractions based on different genome sets.

### Linking MAGs to globally important taxa, genome annotation and functional trait conservation analysis

To assess the coverage of globally important taxa (core and CRAT taxa) within the MiDAS-HQ catalog, 16S rRNA genes extracted from the HQ MAGs were aligned to core and CRAT genera (94.5% identity cutoff) identified in the MiDAS 4 study^10,37^ based on the MiDAS (v4.8) database. Core taxa were first identified separately within each process type (C, CN, CNDN and CNDNP) and subsequently combined to obtain the global core set in the MiDAS 4 stduy^10^. Additionally, core genera were defined as those with a relative abundance of at least 0.1% in each WWTP and were further classified into strict core (>80% of WWTPs), general core (>50%) and loose core (>20%) based on their detection frequencies across WWTPs, as described in the MiDAS 4 study. All HQ MAGs were re-annotated using DRAM (v1.4.6)^104^ with the KEGG database (downloaded in January 2024)^105^. The KEGG-based annotations were then imported into EnrichM (v0.5.0) using the *‘classify’* subcommand (https://github.com/geronimp/enrichM) to identify complete modules for each MAG with the ‘*--custom_modules*’ parameter. The analyzed modules and the criteria for defining complete modules (100% of genes present) followed those established in previous studies^16,106^, the version of the KEGG database used here, or custom modules to construct the functional traits of core genera (**Supplementary Table S12**).

For trait conservation analysis, HQ MAGs assigned to GTDB-defined core genera represented by at least ten genomes were used. For each genus, we calculated the fraction of HQ MAGs encoding each pathway as a measure of within-genus trait conservation. Genus-trait combinations were assigned to three conservation categories: conserved present when the trait was detected in ≥80% of MAGs, mixed when detected in 20-80% of MAGs, and conserved absent when detected in <20% of MAGs. Genera with insufficient genome representation were excluded from the genus-level conservation analysis to reduce the influence of sparse sampling. Then, we extended the trait conservation analysis to higher taxonomic ranks, including family, order, class and phylum, using the same set of HQ MAGs assigned to the 277 core genera. Trait conservation was recalculated independently at each rank following the same category grouping. The fraction of mixed taxon-trait combinations was then compared across ranks to evaluate how functional heterogeneity changed with taxonomic breadth. A follow-up species-level analysis was restricted to traits classified as mixed at the genus level with two cutoffs of at least three and five HQ MAGs per species were evaluated. The resulting species-level mixed fractions were compared with genus-level mixed classifications to assess whether functional heterogeneity within genera could be resolved at the species level.

For nitrifying bacteria, *nxrA* genes were detected with GraftM^107^ using pNXR and cNXR GraftM packages and filtered based on gene phylogeny and gene length of >3000 nucleotides (HMM search lengths: pNXR: 3,411 nucleotides, cNXR: 4,092 nucleotides). Gene-synteny of *nxr*-encoding scaffolds was assessed using the DRAM annotation output in R v.4.5.0 using packages gggenes (v.0.6.0)^108^ and tidyverse (v.2.0.0)^109^. The rTCA-cycle was assessed using DRAM annotations (ko_id: K15230, K15231, K00169, K28726, K28727, K00170, K28728, K00172, K28729, K00171, K01902, K01903, K00031 and kegg_hits: “2- oxoglutarate:ferredoxin oxidoreductase”). The dsrA and dsrB gene pairs from each HQ MAG were concatenated and aligned with reference dissimilatory sulfite reductase (Dsr) sequences. Phylogenetic placements were inferred using IQ-TREE^110^ to determine whether the sequences belonged to the reductive or oxidative Dsr groups.

### Identification and phylogenetic classification of L-*nosZ* genes

L-*nosZ* genes in the MiDAS global genome catalog were identified following the workflow described in the original discovery study^63^. Protein sequences derived from DRAM annotations were screened using hmmsearch (HMMER v3.2.1^111^) against a hidden Markov model (HMM) constructed in the above study. HMM hits were filtered using the following criteria: an e-value threshold of <1×10^-5^, a length range of 400-900 amino acids and the presence of conserved NosZ motifs (CX_2_FCX_3_HXEM, DXHH or GXHH, PHG, GPLH and EPH) as suggested. To further verify the clade assignment, the above candidate sequences were combined with the reference NosZ sequences for phylogenetic inference, i.e., sequence alignment using MAFFT (v7.525)^112^, tree construction using FastTree (2.2.0)^98^. NosZ clades were assigned based on phylogenetic placement relative to the reference sequences, using the most recent common ancestor of each reference clade. Only three HQ genomes representing two species, including one species from the core genus *Desulfobacillus*, were identified.

### Phylogenetic diversity evaluation and novel lineage detection

Phylogenetic diversity was estimated using total branch length, while the additional branch length (phylogenetic gain) quantified the phylogenetic contribution of the MiDAS-HQ catalog. To compute these metrics for each taxonomic lineage, the toolbox, GenomeTreeTk (v0.1.8; https://github.com/donovan-h-parks/GenomeTreeTk)^11^ was applied to a phylogenetic tree constructed from a merged HQ species-representative dataset comprising MiDAS and GTDB (r226), generated using GTDB-Tk (v2.4.1). Additionally, a novel species was defined when a species could not be classified at the species level based on GTDB (r226). To evaluate lineage novelty at higher taxonomic levels, the pipeline, MAG phylogeny (https://github.com/aaronmussig/mag-phylogeny)^15^ was used to assess the relative evolutionary divergence (RED) value based on the entire-genome set of species representatives from GTDB, inferred by PhyloRank (v0.1.12; https://github.com/donovan-h-parks/PhyloRank). Manual review and curation were performed on phylogenies of highly divergent MiDAS-HQ MAGs.

### Naming of novel core taxa

HQ MAGs with high contiguity (≤10 contigs) that remained unclassified in GTDB (r226) among AS core taxa were selected for name proposals under the SeqCode registry (https://registry.seqco.de/), following the previous work^17^. Latinized genus names were derived from the cities where the corresponding WWTPs are located. Genus suffixes were assigned according to genomic features: -*bacter* for representative genomes encoding shape-determining proteins, -*coccus* for genomes lacking shape-determining proteins, -*plasma* for genomes lacking both shape-determining proteins and peptidoglycan biosynthesis, and -*monas* for genomes encoding flagella. Species epithets were assigned using three categories, i.e., generic, WWTP-related, and geographic (country or broader region), to minimize name redundancy and provide geographic representation among proposed names. For lineages whose parent taxa were represented by placeholder names in GTDB, Latinized names were derived from the corresponding genus names. Etymologies for all genus and species names are provided in **Supplementary Table S10**, and the names will be manually registered under SeqCode after publication.

## Data availability

All sequenced nanopore long reads and recovered MAGs (including HQ and MQ) are deposited in ENA under BioProject PRJEB83983. The HQ MAGs belonging to core genera, along with KO assignments, predicted genes, and KEGG-based annotations, are deposited in Zenodo at 10.5281/zenodo.19066579. All MAG statistics are available in the Supplementary Tables. The MiDAS 4.8 database can be accessed at https://www.midasfieldguide.org/guide/downloads. Additional data used in this study are described in the Methods section.

## Competing interests

The authors declare no competing interests.

## Code availability

Code and datasets used to generate the figures are available at https://github.com/Hydro3639/MiDAS-genome.

## Acknowledgment

This work was supported by research grants from the Independent Research Fund Denmark (grant 2035-00360B to M.K.D.D), Novo Nordisk Foundation (REThiNk, grant NNF22OC0071498 to P.H.N and M.K.D.D; grant NNF24OC0095323 to L.L.), Poul Due Jensen Foundation (grant MicroFlora Danica to M.A. and P.H.N.), Villum Foundation (grant 50093 to M.A. and Dark Matter, grant 13351 to P.H.N.) and the European Union (ERC grant 101078234 to M.A.). We also thank the staff of all participating WWTPs for providing samples and assisting in obtaining the necessary permissions for their analysis in this study. Rune Bakke passed away December 15, 2020. David Jenkins passed away March 6, 2021.

## MiDAS Global Consortium

Sonia Arriaga^3^, Rune Bakke^4^, Nico Boon^5^, Mathew Brown^6^, Magnus Christensson^7^, Adeline Seak May Chua^8^, Eddie Cytryn^9^, Leonardo Erijman^10^, Claudia Etchebehere^11^, Despo Fatta-Kassinos^12^, Dominic Frigon^13^, April Z. Gu^14^, Harald Horn^15^, David Jenkins^16^, Norbert Kreuzinger^17^, Ana Lanham^18^, Yingyu Law^19^, TorOve Leiknes^20^, Eberhard Morgenroth^21,22^, Adam Muszyński^23^, Steve Petrovski^24,25^, Maite Pijuan^26^, Suraj Babu Pillai^27^, Maria A. M. Reis^28^, Simona Rossetti^29^, Robert Seviour^30^, Nick Tooker^31^, Pirjo Vainio^32^, Mark C.M. van Loosdrecht^33,1^, Jiří Wanner^34^, David Weissbrodt^33,35^, Tong Zhang^36^ & Per H. Nielsen^1^

^3^Environmental Science Department, The Institute for Scientific and Technological Research of San Luis Potosi (IPICYT), San Luis Potosí, Mexico. ^4^Department of Process, Energy and Environmental Technology, University College of Southeast Norway, Porsgrunn, Norway. ^5^Center for Microbial Ecology and Technology, Ghent University, Ghent, Belgium. ^6^Environmental Engineering, School of Engineering, Newcastle University, Newcastle, England. ^7^Veolia Water Technologies AB, AnoxKaldnes, Lund, Sweden. ^8^Department of Chemical Engineering, Faculty of Engineering, Universiti Malaya, Kuala Lumpur, Malaysia. ^9^The Cytryn Lab, Microbial Agroecology, Volcani Center, Agricultural Research Organization, Rishon Lezion, Israel. ^10^INGEBI-CONICET, University of Buenos Aires, Buenos Aires, Argentina. ^11^Department of Biochemistry and Microbial Genomics, Biological Research Institute “Clemente Estable”, Montevideo, Uruguay. ^12^Department of Civil and Environmental Engineering and Nireas-International Water Research Center, University of Cyprus, Nicosia, Cyprus. ^13^Environmental Engineering, McGill University, Montreal, QC, Canada. ^14^School of Civil and Environmental Engineering, Cornell University, Ithaca, NY, USA. ^15^Water Chemistry and Water Technology and DVGW Research Laboratories, Karlsruhe Institute of Technology (KIT), Karlsruhe, Germany. ^16^David Jenkins & Associates, Inc, Kensington, CA, USA. ^17^Institute for Water Quality and Resource Management, TU Wien, Vienna, Austria. ^18^Water Innovation and Research Centre, University of Bath, Bath, England. ^19^Singapore Centre of Environmental Life Sciences Engineering (SCELSE) Nanyang Technological University, Singapore, Singapore. ^20^Water Desalination and Reuse Center, King Abdullah University of Science and Technology (KAUST), Thuwal, Saudi Arabia. ^21^ETH Zürich, Institute of Environmental Engineering, 8093 Zürich, Switzerland. ^22^ Eawag, Swiss Federal Institute of Aquatic Science and Technology, 8600 Dübendorf, Switzerland. ^23^Faculty of Environmental Engineering, Warsaw University of Technology, Warsaw, Poland. ^24^Department of Microbiology, Anatomy, Physiology and Pharmacology, La Trobe University Bundoora, Australia. ^25^La Trobe Institute for Molecular Sciences, La Trobe University Bundoora, Australia. ^26^Technologies and Evaluation Area, Catalan Institute for Water Research, ICRA, Girona, Spain. ^27^VA Tech Wabag Ltd, Chennai, India. ^28^Biochemical Engineering Group, Universidade Nova de Lisboa, Lisboa, Portugal. ^29^Water Research Institute,National Research Council (CNR-IRSA), Rome, Italy. ^30^La Trobe University, Melbourne, VIC, Australia. ^31^Department of Civil and Environmental Engineering, University of Massachusetts Amherst, Amherst, MA, USA. ^32^Kemira Oyj, Espoo R&D Center, Espo, Finland. ^33^Environmental Biotechnology, TU Delft, Delft, The Netherlands. ^34^Department of Water Technology and Environmental Engineering, University of Chemistry and Technology, Prague, Czech Republic. ^35^Department of Biotechnology and Food Science, Norwegian University of Science and Technology (NTNU), Trondheim, Norway. ^36^Environmental Microbiome Engineering and Biotechnology Laboratory, Department of Civil Engineering, The University of Hong Kong, Hong Kong SAR, China.

## Notes

### Competing Interest Statement

The authors have declared no competing interest.

