## Supplementary Note for "The MiDAS global genome catalog: 53,501 long-read MAGs representing all core prokaryotic genera in the global activated sludge microbiome"

### Microbial diversity in global WWTPs

Activated sludge (AS) communities in global wastewater treatment plants (WWTPs) are estimated to be roughly an order of magnitude less diverse than global ocean microbiomes[1]. Yet they remain among the most complex engineered microbial ecosystems on Earth. From 83 ultra-deep sequenced WWTP samples collected worldwide, we reconstructed 53,501 metagenome-assembled genomes (MAGs), representing 25,221 microbial species. As expected, recovery of high-quality (HQ) MAGs required substantially higher sequencing depth than recovery of medium-quality (MQ) MAGs (median coverage 21 *vs.* 8) (**Fig. 1c**). Despite capturing all core taxa and recruiting ~78% of the total microbial community, the species accumulation curve remained close to linear (**Fig. 2b**). This indicates that many low-abundance taxa persist across samples and that species richness is far from saturation, indicating a long tail of rare species in AS communities[2].

Including MQ MAGs increased the number of known, GTDB-classified lineages detected in global WWTPs to 94 phyla, 232 classes, 635 orders, 1,393 families, 3,585 genera, and 3,170 species (relative to GTDB r226). The remaining species-level clusters (87.4%) lacked GTDB species assignments, consistent with extensive novelty in the MiDAS-HQ-based analysis. Although diversity was high, it was unevenly distributed. The ten most genome-rich phyla accounted for ~89% of recovered MAGs and ~87% of species (**Fig. S11**). This pattern suggests intense selective pressures in WWTPs that favor a limited set of dominant, well-adapted lineages, alongside a diverse background of low-abundance taxa[1,3].

### The MiDAS catalog expands genomic diversity across the tree of life and improves genome quality

We quantified phylogenetic diversity (PD) and phylogenetic gain (PG) of the MiDAS HQ catalog using cumulative branch length, following a previous study[4], in which PD represents the total branch length spanned by a set of taxa, and PG reflects the additional branch length contributed by newly included taxa. A phylogeny tree constructed from 45,267 bacterial HQ species representatives (33,272 from GTDB r226 and 11,995 from MiDAS) showed that MiDAS contributes 39.0% of total PD and increases PG by 26.0% across the bacterial tree represented by HQ genomes (**Fig. S3**). Substantial gains were also observed within the three most genome-rich phyla in GTDB r226 (Pseudomonadota, Actinomycetota, and Bacteroidota), which together account for >68% of GTDB genomic representatives. Within these phyla, MiDAS expanded PD by 19.6-27.0%. Overall, MiDAS increased PD by >20% in 41 bacterial phyla and 102 classes. Take Patescibacteriota as an example, MiDAS contributed 1,311 species and increased the number of HQ species representatives by ~2.5-fold, corresponding to a PG of 47.0% (**Fig. S3** and **Supplementary Table S14**).

Beyond expanding phylogenetic breadth, MiDAS improved genome quality for many previously classified taxa (as measured by CheckM2-based genome completeness). Across 4,187 known taxa spanning 13 phyla, 53 classes, 192 orders, 513 families, 1,710 genera, and 1,706 species, MiDAS provided higher-quality representatives (**Supplementary Table S15**). When comparing MiDAS HQ species representatives with all 113,104 GTDB species representatives, MiDAS had the highest completeness for 6 of the 74 bacterial phyla. When the comparison was restricted to HQ representatives across both datasets, the number increased to 14 bacterial phyla. In addition, MiDAS provides HQ genome representatives for higher-rank lineages that previously lacked HQ genomic representation in GTDB r226, including three phyla (4 species; 5 MAGs), 18 classes (49 species; 59 MAGs), 76 orders (235 species; 312 MAGs), 266 families (968 species; 1,466 MAGs) and 1,265 genera (3,380 species; 5,517 MAGs). Finally, inclusion of MiDAS genomes more than doubled the number of available HQ species representatives for 19 bacterial phyla, 79 classes, 262 orders, 668 families, and 2,021 genera (**Supplementary Table S16**). For example, Bdellovibrionota and Myxococcota increased by 457% and 244%, reaching 490 and 694 HQ species, respectively, of which MiDAS uniquely contributed 402 and 492.

**Genus-level taxonomy provides a reliable proxy for functional potential**

The high trait conservation (91.6%, including conserved present and absent) suggested the usefulness of genus as a useful taxonomic unit to profile the functional potential of the AS community. While trait conservation also differed among functional categories (**Fig. S7a and b**). Central carbon and energy metabolism provided the expected shared metabolic backbone, while nitrification represented a highly restricted and genus-conserved specialized guild. By contrast, complete denitrification was conserved in only seven core genera and mixed in 21, consistent with modular distribution of nitrogen-reduction steps across taxa. Intracellular storage polymers and sulfur-oxidation traits showed intermediate patterns, with some modules conserved in subsets of core genera, while others were mixed or rarely recovered, highlighting the need for genome-resolved interpretation.

To further determine whether genus-level mixed patterns reflected finer-scale species differences, we performed a species-level follow-up restricted to pathways that were mixed at the genus level and to genome-defined species represented by at least three or five HQ MAGs. Mixed fractions decreased from 8.4% at the genus level to 5.9% and 4.3% in the ≥3- and ≥5-MAG species-level follow-up analyses, respectively (**Fig. S7c**). Although this comparison is limited to species with sufficient genome representation, the reduction in mixed classifications suggests that some genus-level functional heterogeneity is resolved at the species level. Together, these results show that genus-level taxonomy provides a useful unit for interpreting many AS traits, especially specialized guilds such as nitrifiers, but that modular or annotation-sensitive traits such as denitrification, carbon fixation and storage metabolism require species- or genome-resolved interpretation.

**Metabolic potential differentiation of polyphosphate-accumulating organisms**

Polyphosphate-accumulating organisms (PAOs) are a group of bacteria used in the enhanced biological phosphorus removal (EBPR) technology to capture phosphorus (P) from wastewater. These bacteria are able to store large amounts of inorganic P intracellularly as polyphosphate and cycle it through anaerobic-oxic feast-famine phases. Currently, four bacterial genera comprise the confirmed PAO species: *Ca.* Accumulibacter, *Azonexus* (formerly *Dechloromonas*), *Ca.* Phosphoribacter and *Ca.* Lutibacillus (both formerly *Tetrasphaera*). *Ca.* Accumulibacter and *Azonexus* generally also contain PHA and glycogen, while the content of these storage polymers in *Ca.* Phosphoribacter and *Ca.* Lutibacillus is more uncertain.

Comparative genomic profiling revealed clear lineage-specific metabolic patterns within individual genera. For polyphosphate (poly-P) accumulation, nearly all species encoded the core *pit* and *pstSCAB*-*phoU* transport systems (100%, 5/5 in *Ca.* Lutibacillus; 98%, 39/40 in *Ca.* Accumulibacter; 89%, 25/28 in *Ca.*

Phosphoribacter; 85%, 44/52 in *Azonexus*), supporting a widespread capacity for high-affinity phosphate import (**Fig. 6**). In contrast, the associated regulatory components (*phoB*-*phoR)*[5] showed clear lineage-specific differences. The gene set was highly conserved in *Ca.* Accumulibacter (>95%) and *Azonexus* (85%), but absent from *Ca.* Phosphoribacter and *Ca.* Lutibacillus, indicating that phosphate transporter control in the latter genera may rely on alternative or as-yet-unknown regulatory mechanisms[6,7].

Several glycogen-associated genes (*glgB*, *glgC*, *glgU,* and *glgP*) were widely conserved (78% of species) across the four genera. Yet, the key *glgA* gene for glycogen synthesis was detected in most *Ca.* Accumulibacter (95%, 38/40) and *Azonexus* (65%, 34/52) species, but in none of the *Ca.* Phosphoribacter or *Ca.* Lutibacillus species, consistent with the lack of in situ glycogen detection in these lineages[6]. A similar pattern was observed for PHA synthesis, with the *phaABC* genes nearly universal in *Ca.* Accumulibacter (100%) and *Azonexus* (98%), but less prevalent in *Ca.* Phosphoribacter (*phaB*, 14%; *phaC*, 39%) and completely absent (*phaC*) in *Ca.* Lutibacillus, suggesting a shift toward alternative carbon-storage strategies.

Cyanophycin has been proposed as a potential storage polymer for *Ca.* Lutibacillus and *Ca.* Phosphoribacter[6]. Although cyanophycin storage is not restricted to PAOs, consistent with and extending this view, the biosynthetic gene *cphA* was detected in nearly all *Ca.* Accumulibacter, *Ca.* Lutibacillus*,* and *Azonexus* species (98-100%), but in only half of *Ca.* Phosphoribacter species (16/28). Conversely, the degradative *cphB* was fully conserved in *Ca.* Lutibacillus (5/5) but rare in *Ca.* Accumulibacter (1/40) and *Azonexus* (5/52), with *Ca.* Phosphoribacter intermediate (16/28). This observation suggests that cyanophycin-based storage and its subsequent mobilization are likely particularly central to *Ca.* Lutibacillus, whereas *Ca.* Accumulibacter and *Azonexus* may use cyanophycin in a more facultative manner.

Denitrification traits also showed strong lineage specificity. *Ca.* Azonexus (40/52) was the only genus in which a complete denitrification pathway was common, consistent with a potential dual role in nitrogen and phosphorus removal[7]. *Ca.* Accumulibacter was dominated by *nirS* (93%, 37/40) and retained *NosZ* in about half of its species (53%, 21/40). In contrast, *Ca.* Phosphoribacter and *Ca.* Lutibacillus mainly encoded the alternative *NirK*-type nitrite reductase (54% and 40%, respectively) but lacked the *NosZ* gene, implying truncated denitrification chains. Beyond nitrite reductase composition, the four PAO genera also differed in their nitrate-reductase systems. *Ca.* Accumulibacter species displayed a mutually exclusive distribution of the periplasmic *NapAB* complex (21/40 species), which is typically expressed under more oxic or fluctuating conditions[8–10] and the membrane-bound *NarGHI* complex (18/40 species), which supports more energy-efficient nitrate respiration under low-oxygen conditions[11]. *Ca.* Azonexus predominantly encoded *NapAB* (51/52), whereas *Ca.* Phosphoribacter (13/28) and *Ca.* Lutibacillus (3/5) favored the periplasmic *NarGHI.* Together, these results indicate that the four PAO genera span a continuum from fully denitrifying, *Nap*-based PAOs (*Azonexus*) to *Nar*-dominated, truncated denitrifiers (*Ca.* Phosphoribacter and *Ca.* Lutibacillus), with *Ca.* Accumulibacter occupying an intermediate, flexible niche. As a consequence, the PAO genera differ in how they couple phosphorus removal to specific nitrogen oxidation and reduction pathways along redox gradients.

Overall, beyond phosphate removal, *Azonexus* emerges as a *Nap*-based complete denitrifier, suggesting that it is particularly suited to micro-oxic and mildly anoxic zones where phosphate uptake can be tightly coupled to nitrate reduction[7]. As partial denitrifiers, *Ca.* Accumulibacter spans a broader niche, with coexisting Nap- and Nar-based lineages, suggesting a flexible PAO strategy operating under fluctuating redox conditions. In contrast, *Ca.* Phosphoribacter and *Ca.* Lutibacillus appear as *Nar*-dominated, *NirK*-type PAOs, traits that are more compatible with strongly anoxic, nitrate-rich microenvironments, such as deeper floc or biofilm regions. Taken together, these complementary PAO strategies suggest that EBPR communities partition phosphorus-removal functions along redox gradients, which may help stabilize process performance under variable loading and aeration regimes.

**The discrepancy between 16S rRNA gene-based and genome-based classification**

Through an extensive, targeted sequencing effort of AS metagenomes from full-scale WWTPs worldwide, we constructed the MiDAS global genome catalog, comprising 53,501 highly contiguous MAGs clustered into 25,221 microbial species, including 12,047 prokaryotic species. This catalog substantially expands the known phylogenetic diversity of AS microbial communities and provides the first HQ genome representatives for large portions of both previously named and novel taxa across multiple deep ranks. It thereby brings previously unrepresented branches of the WWTP microbial tree of life into genomic view (**Fig. S3 and Supplementary Table S7**). High read recruitment across both the 83 metagenomes generated here (median 77.6%) and 295 external AS metagenomes (median 72.8%) indicates that the catalog captures most dominant populations and a substantial abundance-weighted fraction of genomic diversity in global AS microbiomes. By linking genomes to the MiDAS gene-based taxonomy[3] *via* their 16S rRNA genes, which are recovered in ~85% of MAGs, this resource establishes a unified reference framework that connects genome-resolved analyses with long-term 16S rRNA gene-based microbial surveillance of wastewater treatment systems[12]. Here, focusing on the collection of HQ MAGs, we evaluated concordance between 16S rRNA gene-based and genome-based taxonomic classification. We classified 16S rRNA genes against the ecosystem-specific MiDAS 5.3 database and classified genomes using GTDB. As expected, genome-based classification showed lower species-level assignment rates, reflecting extensive novelty among MiDAS genomes relative to GTDB. By contrast, 16S rRNA gene classification achieved higher apparent species-level resolution; however, 95% of assigned "species" carried *de novo* MiDAS identifiers (for example, *midas_s_2783*), highlighting the limited linkage of 16S rRNA gene-based labels to genomically defined species.

For both classification methods, assignment success increased toward higher taxonomic ranks. Discrepancies were most significant at the species rank and decreased at higher ranks, except for the phylum level. At this level, differences were dominated by nomenclature, most notably MiDAS "Proteobacteria" corresponding to GTDB "Pseudomonadota" (**Supplementary Table S17**), reflecting recent revisions to prokaryotic taxonomy and nomenclature[13]. Overall, 16S rRNA gene- and genome-based classifications were broadly consistent at higher ranks but diverged substantially at lower ranks, mainly due to the novelty of the recovered MiDAS species. These inconsistencies should diminish when the MiDAS genome catalog is incorporated into future GTDB releases and as MiDAS amplicon taxonomy increasingly adopts GTDB as a backbone.

We further examined 10,129 genome-derived species clusters for which 16S rRNA genes could be classified at the species level. These clusters mapped to 8,430 distinct MiDAS 5.3 species names, indicating that multiple genome-defined species often collapse into a single 16S rRNA gene-defined species. For example, 14 genome-defined species were assigned to the same species, *Nitrospira defluvii*, based on 16S rRNA gene classification. This collapse highlights a key limitation of 16S rRNA genes in resolving finescale diversity and suggests that 16S rRNA gene-based taxonomy can underestimate species richness compared with genome-based approaches. Such mismatches also pinpoint high-priority taxa for revising ecosystem-specific taxonomy and for uncovering novel diversity through the genome-resolved approach[6].

### Supplementary Figures

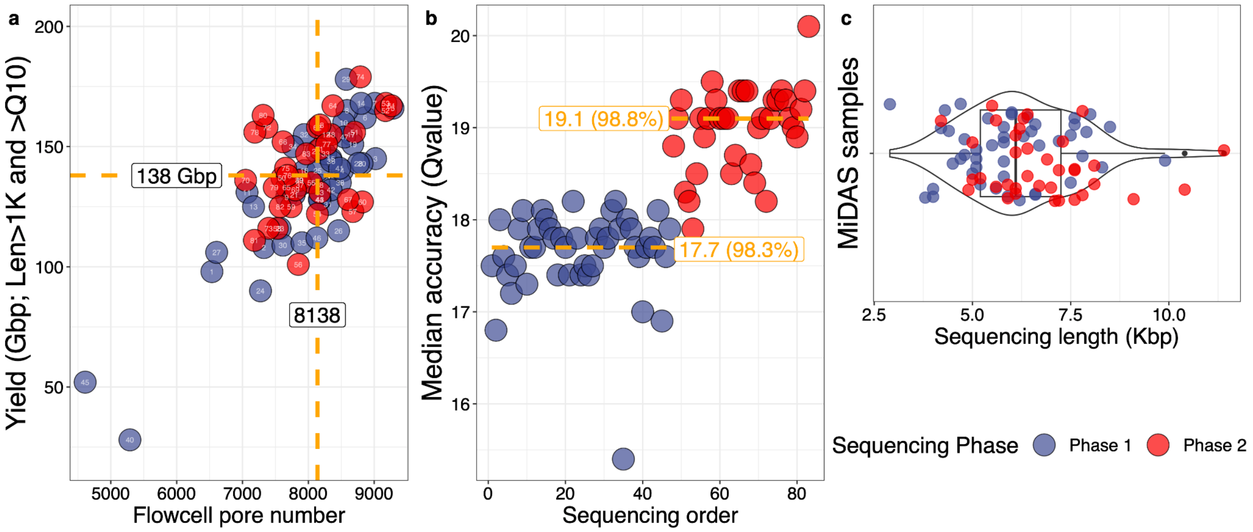


Figure S1 | Sequencing statistics of MiDAS samples. a, sequencing yield, b, read accuracy, and c, read-length distribution for each sample, based on trimmed nanopore reads after filtering for lengths ≥1 kb.

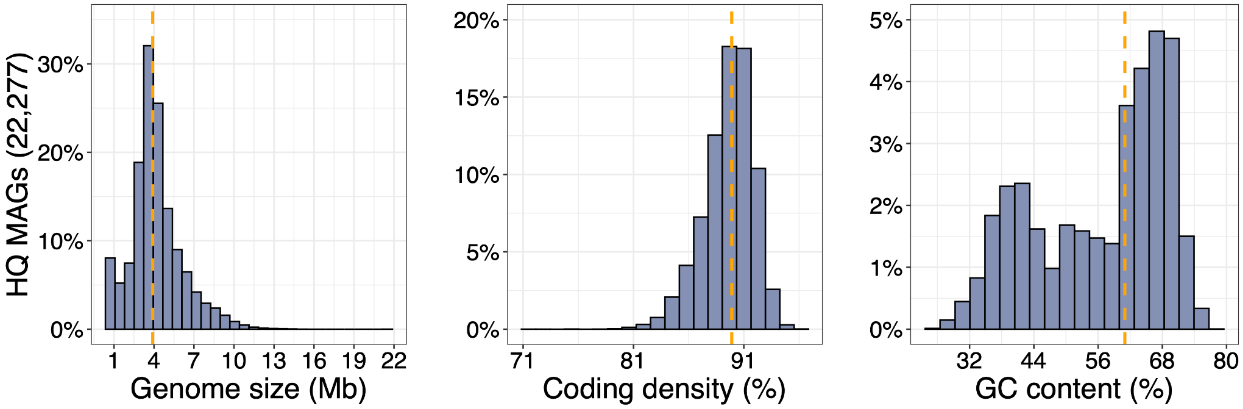


Figure S2 | Genomic statistics of high-quality MAGs in the MiDAS catalog.

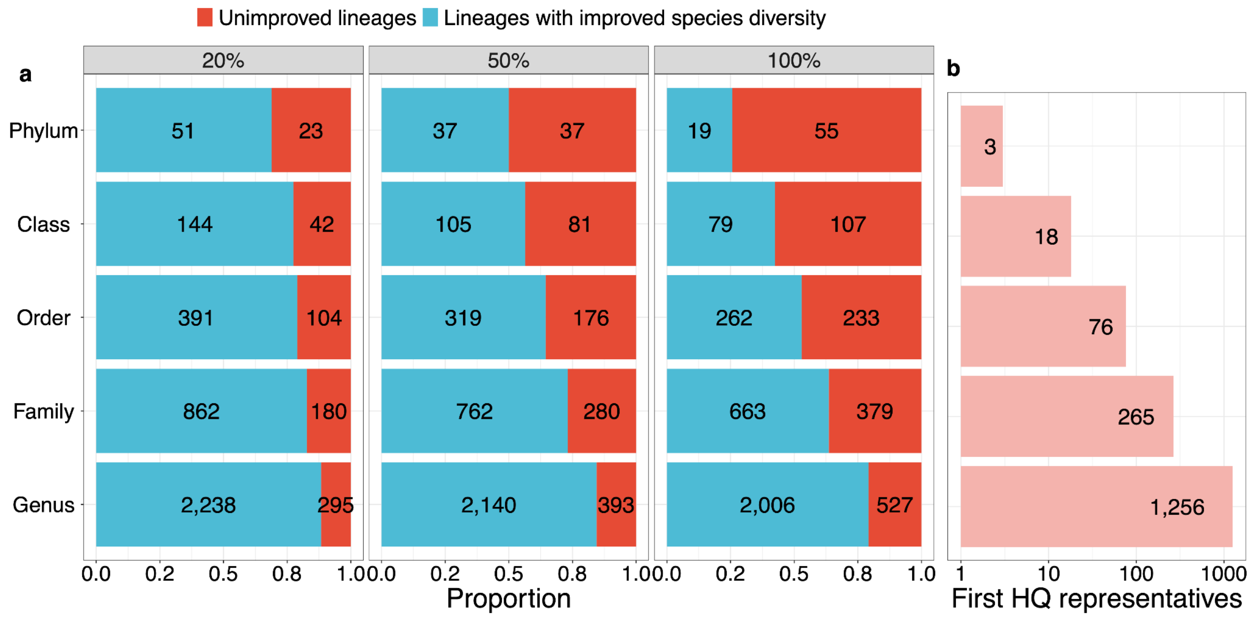


Figure S3 | Increased bacterial species diversity and first HQ representatives contributed by MiDAS HQ relative to GTDB r226. (a) Number of known taxa (by rank) for which inclusion of MiDAS HQ species representatives increases species diversity by at least 20%, 50%, or 100% relative to GTDB r226 (three columns). Genera and higher taxa lacking HQ MAG representatives in GTDB r226 were assigned to improvement categories after incorporation of their first HQ MAG introduced by MiDAS. (b) Number of known taxa for which MiDAS provides the first HQ genome representatives.

GTDB PD
MiDAS PD
MiDAS PG/PD

| | HQ species representatives | | | | |
|---|---|---|---|---|---|
| | GTDB.R226 | MiDAS | Total PD | Relative Taxon PD | MiDAS PG |
| **Bacteria** | 33,271 | 11,995 | 100.0% | | 26.0% |
| ARS69 | 0 | 2 | 0.0% | | 100.0% |
| Acidobacteriota | 492 | 425 | 2.6% | | 27.0% |
| Actinomycetota | 6,285 | 1,128 | 9.5% | | 19.6% |
| Babelota | 48 | 43 | 0.6% | | 35.9% |
| Bacteroidota | 3,840 | 1,856 | 9.8% | | 27.0% |
| Bdellovibrionota | 87 | 372 | 1.8% | | 71.8% |
| Bdellovibrionota_G | 2 | 27 | 0.1% | | 80.8% |
| CLD3 | 3 | 7 | 0.0% | | 45.2% |
| Chlamydiota | 83 | 105 | 0.9% | | 49.2% |
| Chloroflexota | 583 | 705 | 4.1% | | 34.1% |
| Cloacimonadota | 6 | 7 | 0.1% | | 30.2% |
| Desulfobacterota_D | 3 | 4 | 0.0% | | 25.7% |
| Desulfobacterota_G | 9 | 9 | 0.1% | | 41.6% |
| Eisenbacteria | 11 | 27 | 0.1% | | 36.5% |
| Electryoneota | 6 | 9 | 0.0% | | 36.8% |
| Elusimicrobiota | 35 | 46 | 0.3% | | 31.0% |
| Fibrobacterota | 18 | 17 | 0.1% | | 28.1% |
| Gemmatimonadota | 119 | 80 | 0.6% | | 19.5% |
| Hydrogenedentota | 15 | 36 | 0.2% | | 50.1% |
| J088 | 0 | 2 | 0.0% | | 100.0% |
| JADJOY01 | 1 | 7 | 0.0% | | 64.6% |
| Krumholzibacteriota | 11 | 14 | 0.1% | | 28.5% |
| Margulisbacteria | 0 | 3 | 0.0% | | 100.0% |
| Myxococcota | 236 | 543 | 2.3% | | 54.5% |
| Patescibacteriota | 808 | 1,311 | 11.4% | | 47.0% |
| Planctomycetota | 413 | 692 | 4.1% | | 46.6% |
| Pseudomonadota | 12,609 | 3,163 | 23.9% | | 19.6% |
| Spirochaetota | 294 | 86 | 1.2% | | 21.7% |
| UBA10199 | 13 | 23 | 0.2% | | 54.4% |
| Verrucomicrobiota | 368 | 451 | 2.5% | | 32.5% |
| Vulcanimicrobiota | 3 | 23 | 0.1% | | 62.8% |
| Zixibacteria | 40 | 22 | 0.3% | | 16.9% |

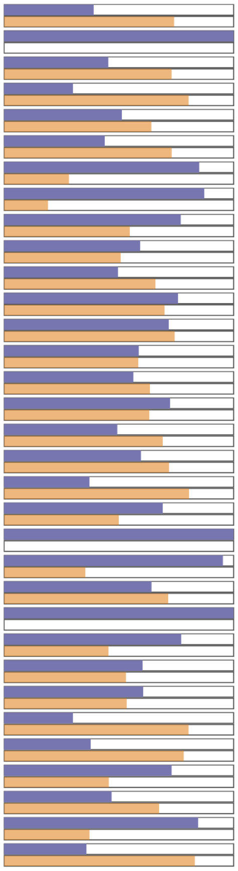

Figure S4 | Phylogenetic diversity (PD) and phylogenetic gain (PG) contributed by MiDAS HQ genomes across selected bacterial phyla. HQ species representatives collected in MiDAS and GTDB r226 were included (MQ genomes were not considered). PD and PG were calculated on a bacterial phylogeny tree inferred from 120 conserved marker genes. Left, number of HQ species per lineage (bacterial domain and individual phyla). Bars show total PD per lineage (normalized to the bacterial domain), the PD contributions of GTDB and MiDAS genomes, and the additional PG contributed by MiDAS genomes.

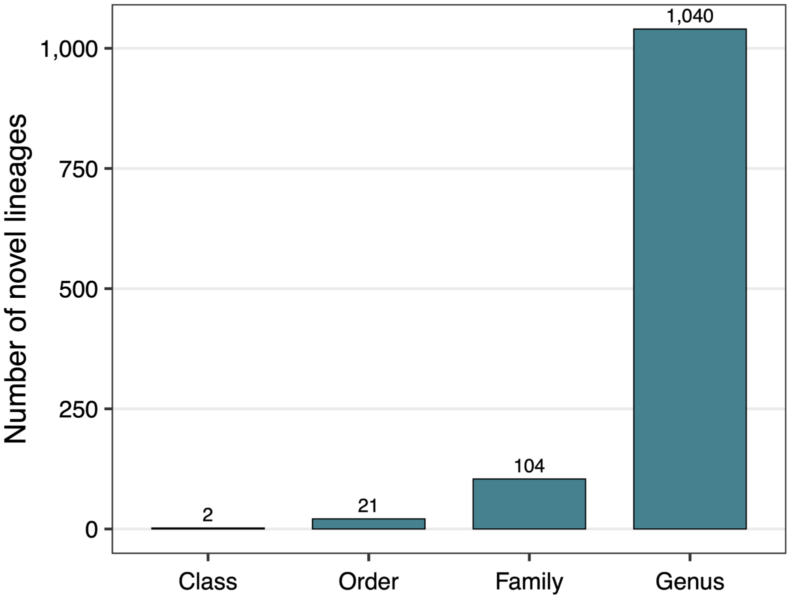


Figure S5 | Distribution of novel lineages inferred from relative evolutionary divergence (RED). RED values for MiDAS HQ lineages are compared with those from the full GTDB r226 dataset.

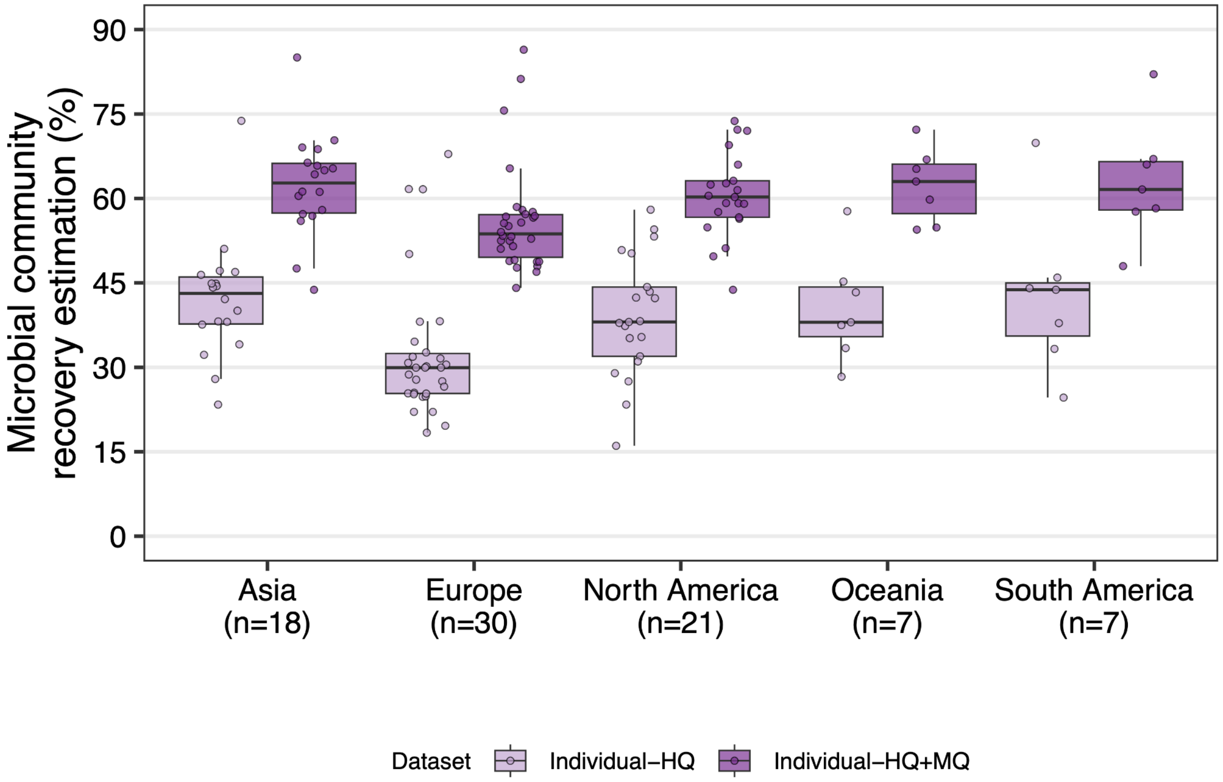


Figure S6 | Microbial community recovery across 83 metagenomes sequenced in this study, estimated using HQ and combined HQ+MQ MAG datasets recovered from individual samples. Recovery was calculated based on the proportion of reads mapped to MAGs using minimap2.

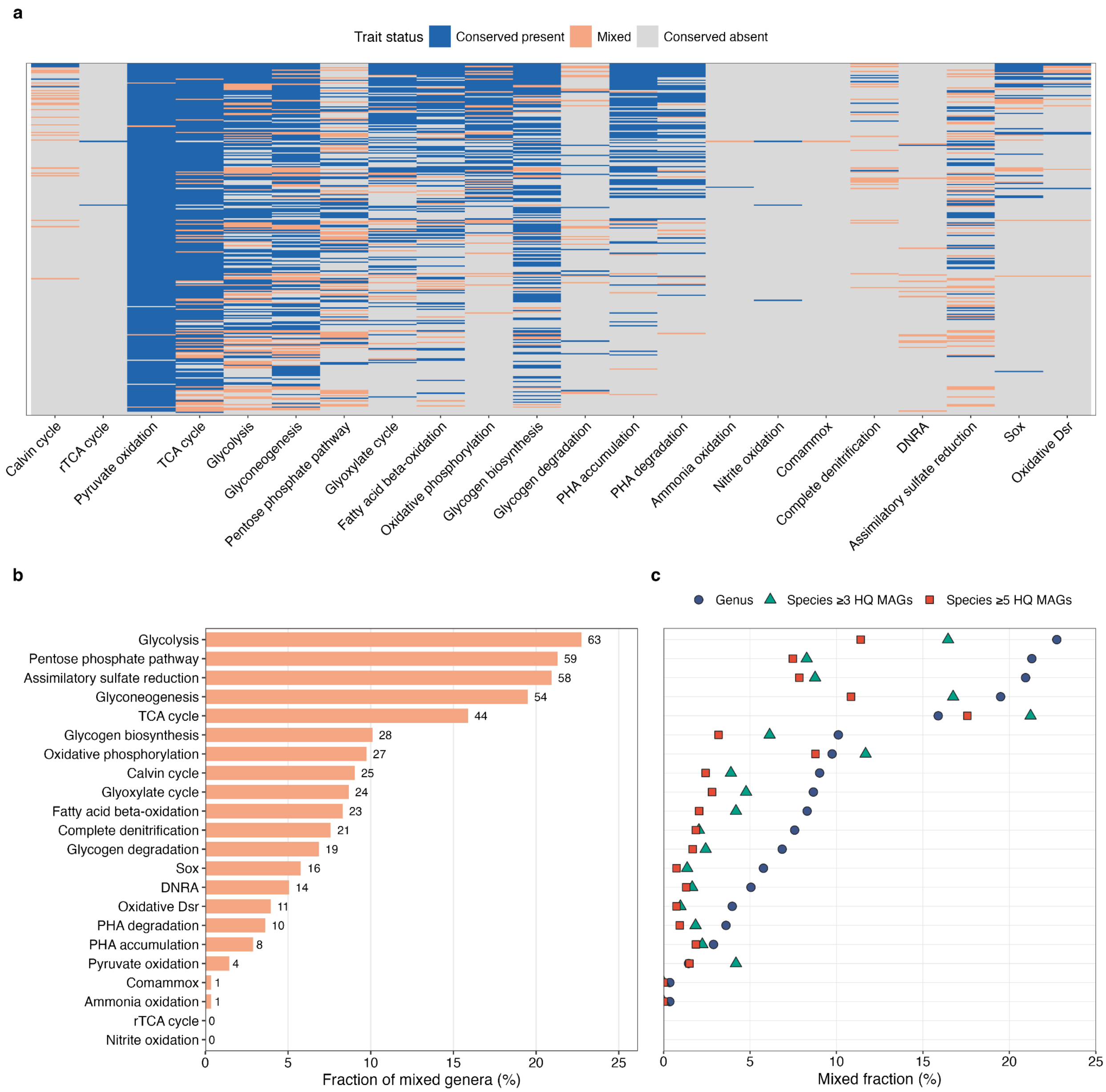


Figure S7 | Genus-level conservation of genome-encoded traits across 277 activated sludge core taxa. a, Conservation status of genome-encoded traits across 277 core genera represented by at least ten HQ MAGs. Traits were classified as conserved present, mixed or conserved absent according to the fraction of MAGs encoding each complete trait within a genus. Each row indicates one core genus and each column represents one inferred functional trait. b, Fraction of core genera classified as mixed for each pathway, showing pathway-dependent variation in genus-level functional consistency. c, Species-level follow-up analysis of traits that were mixed at genus level. Mixed fractions were recalculated for genome-defined species represented by at least three or five HQ MAGs.

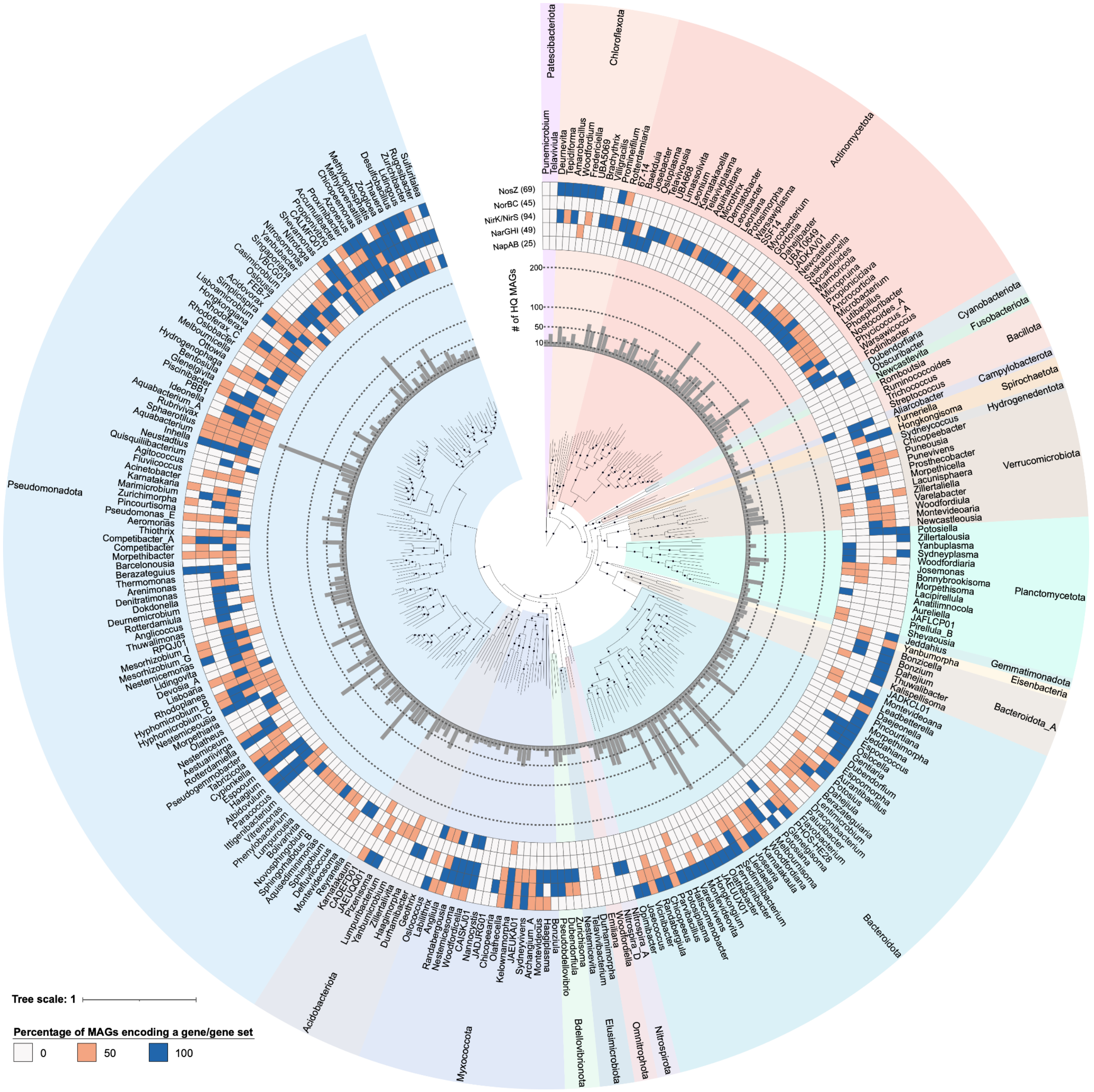


Figure S8 | Denitrification gene profiles across core genera. Inner rings show core genera colored by phylum, followed by the number of HQ MAGs per genus. Heatmap rings show, for each gene, the percentage of HQ MAGs within each genus that encode the gene or gene cluster.

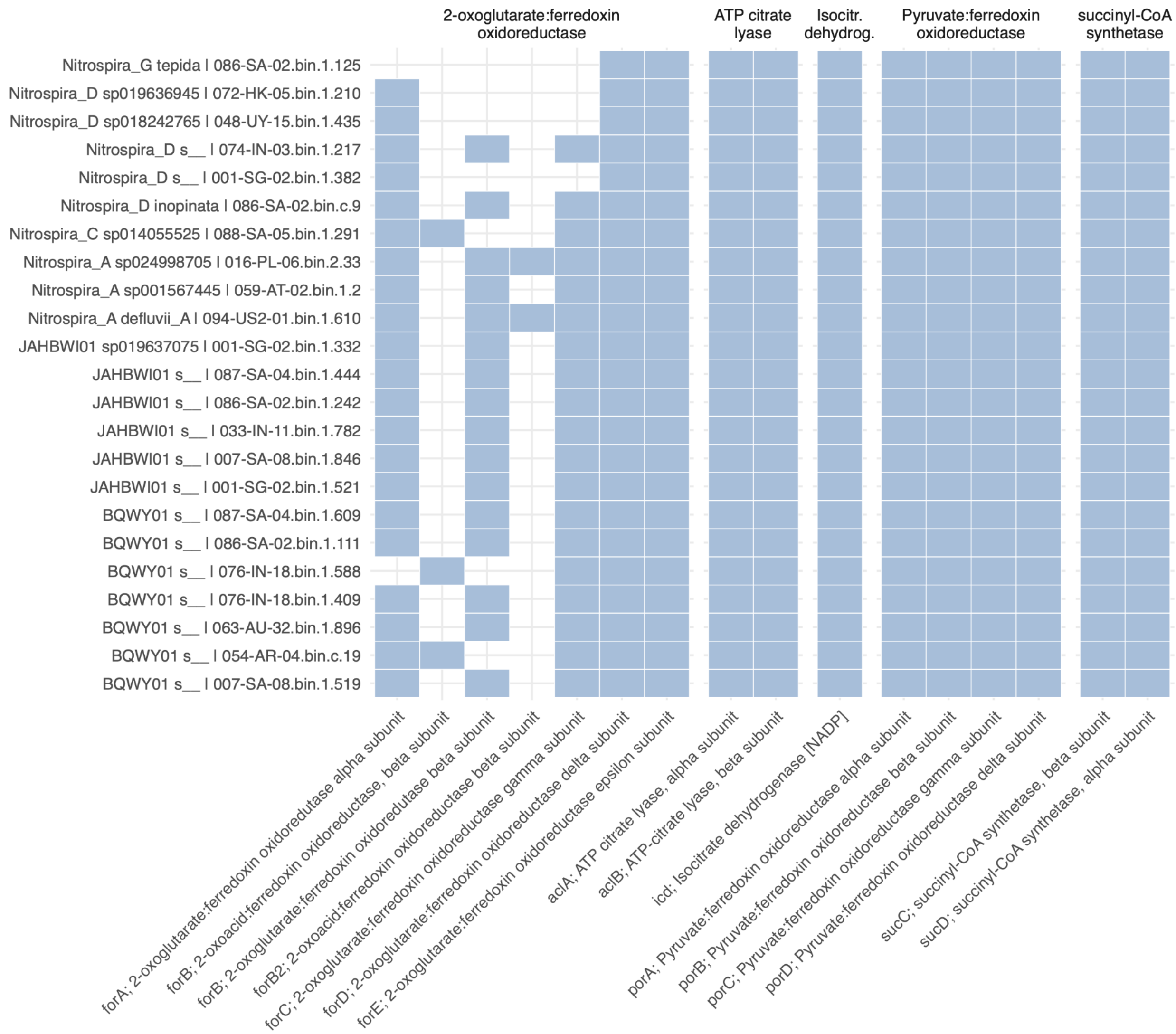


Figure S9 | The presence of key genes and enzymes associated with the reductive tricarboxylic acid (rTCA) cycle identified from all HQ MAGs of the newly identified novel nitrite-oxidizing bacteria and representative HQ MAGs of the canonical NOB genera. The labels on the left show the GTDB-based taxonomic classification and the genome identifier.

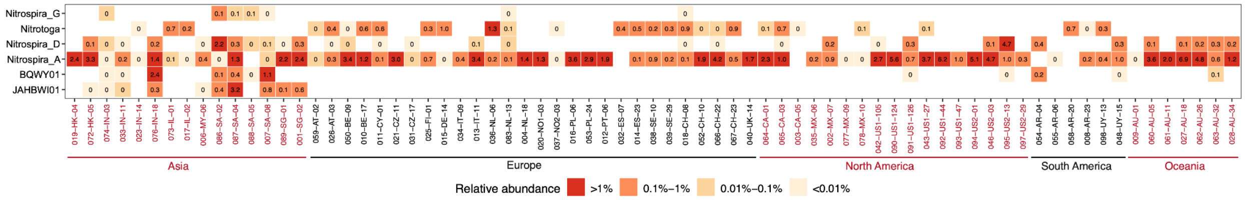


Figure S10 | Global distribution of canonical and newly identified NOB genera across WWTPs. Numbers in cells indicate the cumulative relative abundance of individual species within each genus in each WWTP. WWTPs are grouped by continent, indicated by alternating colors. Sample names include three parts: the first three digits indicate the sequencing order, the middle country code indicates the country where the sample was collected (see **Supplementary Table S1**), and the final number is the sample identifier. For countries with multiple sample coordinators, an additional numeric suffix is added to the country code to distinguish samples coordinated by different groups.

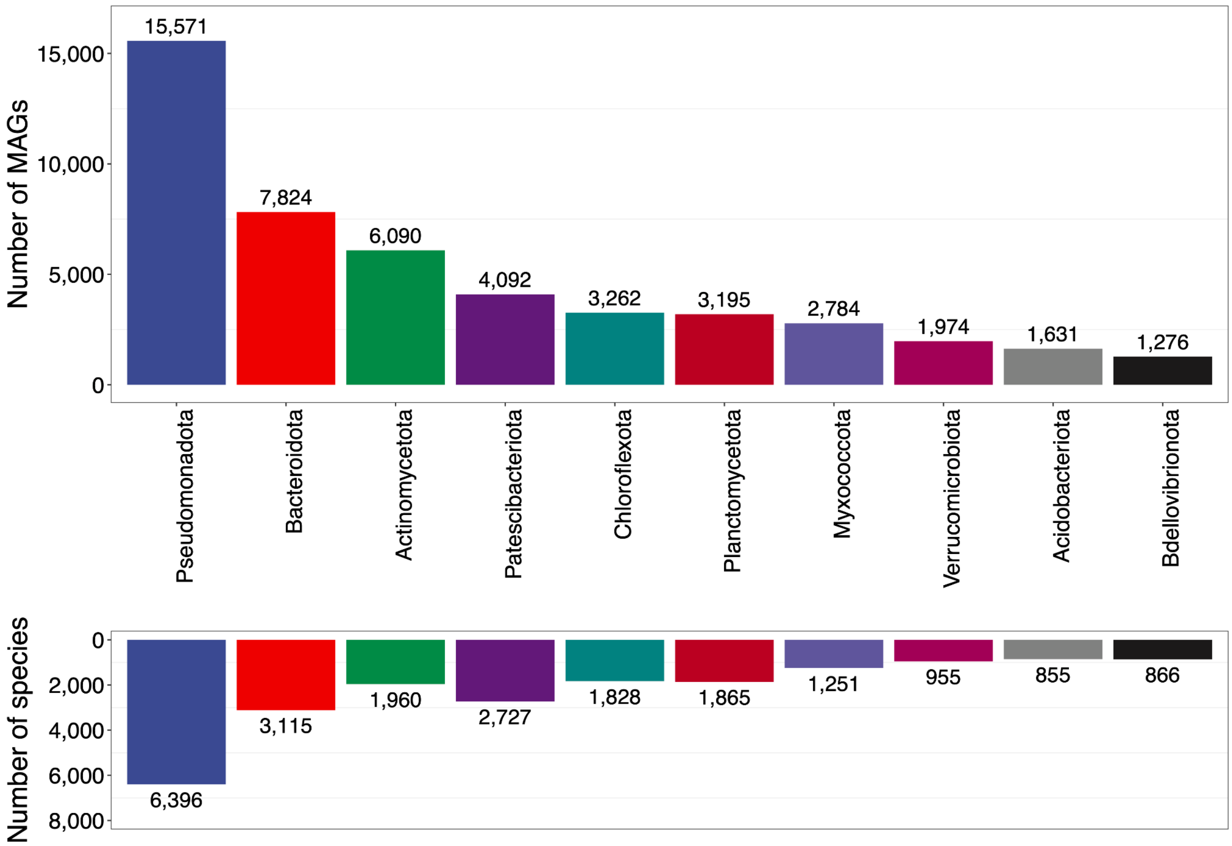


Figure S11 | Ten phyla with the largest numbers of recovered MAGs and species in the MiDAS catalog. Both HQ and MQ MAGs are included.
